# D-gluconate drives *Salmonella* growth during acute and chronic infection

**DOI:** 10.64898/2026.04.07.716905

**Authors:** Christopher Schubert, Melanie Hoos, Andreas Sichert, Nicolas Näpflin, Sanne Kroon, Samuele Pulli, Jeongmin Kim, Lukas Burkhardt, Bidong D. Nguyen, Christian von Mering, Uwe Sauer, Wolf-Dietrich Hardt

## Abstract

Monosaccharides support *Salmonella enterica* serovar Typhimurium colonization of the gut, yet the role of their oxidized derivatives remains understudied. Sugar acids are largely diet-independent carbon sources generated by host-driven oxidative processes, but their contribution during infection - particularly that of less oxidized aldonic and uronic acids - has not been defined. Here, we systematically assess the role of sugar acids derived from D-glucose and D-galactose in *S*. Typhimurium SL1344 colonization. Among D-glucose-derived acids, D-gluconate accumulated to the highest levels and was the dominant substrate supporting luminal expansion in streptomycin-pretreated mice, exceeding the more oxidized acids D-glucuronate and D-glucarate. During chronic infection, D-glucose-derived sugar acids became increasingly important for pathogen persistence. Ecological niche invasion assays identified these compounds as a principal metabolic niche, whereas D-galactose-derived acids contributed minimally. Consistent with a transient, inflammation-linked nutrient niche, sugar acid utilization pathways were similarly prevalent in *Escherichia coli* from individuals with and without inflammatory bowel disease. Together, these findings identify D-gluconate as a key inflammation-dependent nutrient source that fuels *Enterobacteriaceae* expansion in the inflamed gut.

## Introduction

The gut microbiota provides colonization resistance against invading pathogens through two principal mechanisms: interference and exploitation. Interference involves direct antagonism, such as the use of type VI secretion systems, bacteriocins, or contact-dependent inhibition systems. In contrast, exploitation represents indirect antagonism, where commensals compete with pathogens for shared nutrient resources ^1–3^. In recent years, exploitation has been recognized as a key driver of both inter-and intraspecies competition during gut colonization ^4–7^. Despite these defenses, *Salmonella enterica* serovar Typhimurium (*S.* Typhimurium), a Gram-negative enteric pathogen, is capable of overcoming colonization resistance, leading to foodborne diarrheal disease in humans. Animal infection models have revealed that *S.* Typhimurium colonizes the gut in distinct phases ^8^. After oral infection, the pathogen initially expands in the intestinal lumen without disrupting the resident microbiota ^9–11^. This is followed by invasion of the intestinal epithelium, which elicits host immune responses that reduce tissue-associated bacterial loads and reshape the gut luminal environment. During inflammation, *S.* Typhimurium is able to thrive by exploiting inorganic electron acceptors through anaerobic and microaerophilic respiration ^12–15^, whereas L-aspartate–driven fumarate respiration is important in both inflammatory and non-inflammatory phases ^11,16–21^. In contrast, electron donors and carbon sources utilized *in vivo* have only recently been characterized in greater detail. Monosaccharides are key drivers of *S*. Typhimurium colonization, particularly D-glucose, D-fructose, and D-mannose, while D-galactose, fructose-lysine, and D-galactitol play a context-dependent role ^16,18,22–25^. Furthermore, diet-derived L-arabinose released from the polysaccharide arabinan by *S.* Typhimurium promotes pathogen expansion in long-term infection models ^26^. Among these, D-glucose is overall the predominant dietary monosaccharide, accounting for approximately 83.4% of total intake, whereas D-galactose contributes around 4.7% in a representative US cohort ^27^. This aligns with dietary composition data indicating D-glucose as the most abundant monomer in human diet ^28^.

Sugar acids represent an underexplored class of gut metabolites, many of which arise from host-driven oxidation of dietary monosaccharides ^29^. In hexoses such as D-glucose, oxidation can occur at different carbon positions and leads to the formation of distinct classes of sugar acids. Oxidation at C1 converts the aldehyde group (-CHO) into a carboxylate group (-COO⁻), generating an aldonic acid, such as D-gluconate. In contrast, oxidation at C6 targets the primary alcohol group (-CH₂OH) and converts it into a carboxylate group (-COO⁻), producing a uronic acid, such as D-glucuronate. When oxidation occurs at both C1 and C6, the molecule forms a dicarboxylic acid (aldaric acid), exemplified by D-glucarate. Formation of D-glucarate requires stronger or more extensive oxidation than D-gluconate because both the C1 aldehyde and the C6 primary alcohol must be oxidized to carboxylate groups. At the near-neutral pH of the gut (≈ 6-7.5), sugar acids such as D-gluconic acid are predominantly deprotonated to their anionic form, D-gluconate, since their carboxyl groups have p*K*_a_ values of about 3-4. In this anionic form, they are chemically stable but poorly membrane-permeable, requiring dedicated transport systems for uptake ^30^. For example, *S.* Typhimurium ATCC14028s was shown to exploit D-galactarate and D-glucarate for expansion in the inflamed gut following antibiotic treatment. These compounds are generated in the gut lumen through oxidation mediated by iNOS-derived reactive nitrogen species, particularly nitric oxide (NO), a strong oxidant ^29^. NOS2 is strongly upregulated and elevated 4- to 12-fold in patients with active inflammatory bowel disease (IBD) ^31–33^, suggesting that sugar acids may represent an important metabolic niche in chronically inflamed patients. Recent findings support this hypothesis: inflammatory bowel disease (IBD) promotes the evolution and utilization of aldaric acids, particularly D-glucarate and D-galactarate, among enteric pathogens such as *Salmonella* as well as commensal bacteria ^34^, highlighting their ecological and pathogenic relevance. This is supported by the observation that sugar acids are underrepresented among dietary carbohydrates ^27,28^, highlighting host-mediated oxidation as the primary driver of sugar acid formation in the gut.

Since D-glucose and D-galactose are abundant in the murine gut ^11,16,18^, their corresponding sugar acid derivatives are likely readily available and might represent a niche promoting *Salmonella* gut colonization. However, apart from D-glucarate and D-galactarate utilization by S. Typhimurium ATCC14028s ^29^, this hypothesis remains largely untested. It is still unclear which sugar acids are most critical. Given that *S*. Typhimurium encodes the necessary catabolic pathways to utilize them ^35^, we sought to determine which sugar acid most strongly promotes *S*. Typhimurium colonization during early growth and prolonged infection. To this end, we systematically investigated the degradation pathways of the aldonic, uronic, and aldaric acid derivatives of D-glucose and D-galactose in *S*. Typhimurium (**Fig. 1b, and Fig. 2a**). Across a range of murine infection and colonization models, we identified D-gluconate as the most critical sugar acid.

**Figure 1.**
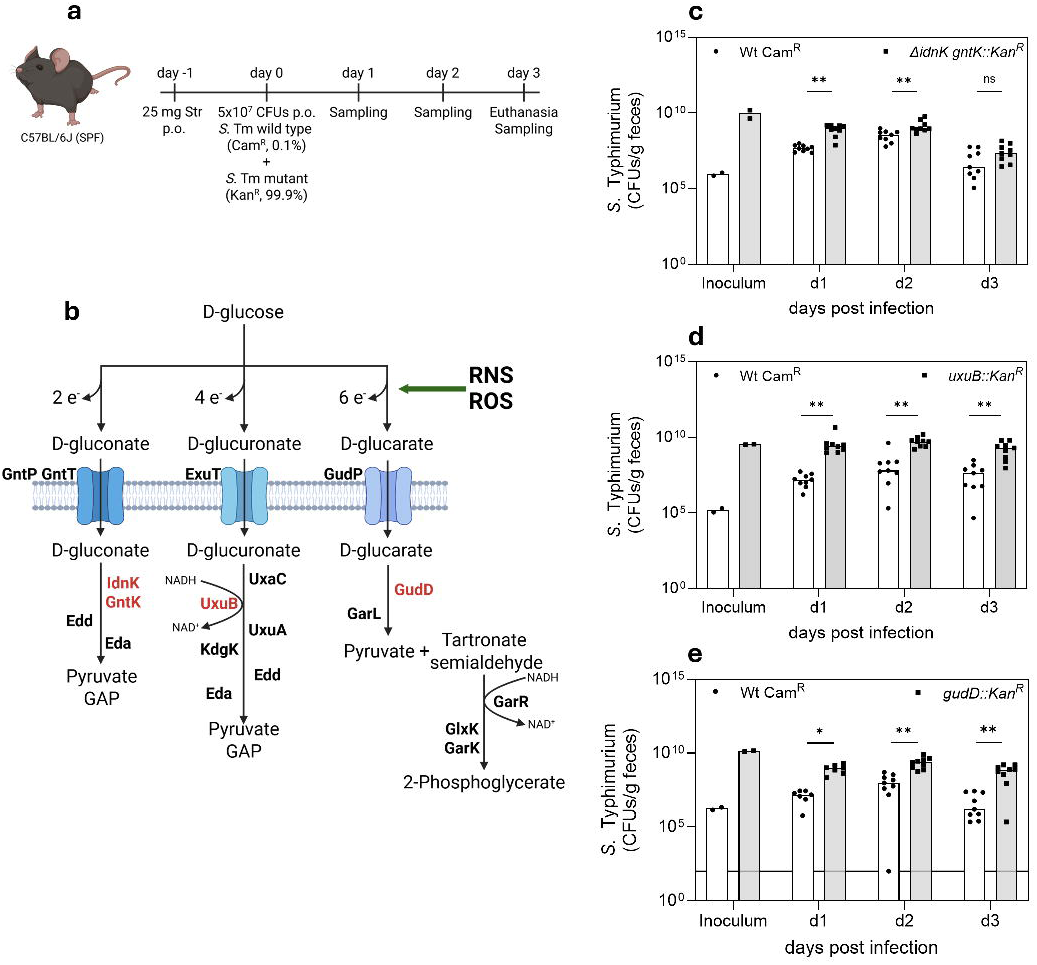
D-gluconate is the primary D-glucose-derived sugar acid driving wild-type expansion. **a**, Experimental design of the streptomycin-pretreated mouse infection model. C57BL/6J specific-pathogen-free mice were treated with streptomycin (25 mg, p.o.) one day before infection and orally inoculated with 5 × 10⁷ CFU of a mixed *S.* Typhimurium wild type and mutant inoculum at a 0.001:1 ratio. Fecal samples were collected daily, and animals were euthanized on day 3 post infection for final sampling. Created in BioRender. Schubert, C. (2026) https://BioRender.com/3p40qc2. **b,** Schematic overview of three D-glucose-derived sugar acid utilization pathways, including D-gluconate, D-glucuronate, and D-glucarate, and their proposed generation under reactive oxygen and nitrogen species (ROS/RNS). Transporters and metabolic enzymes are indicated, and genes targeted in the mutant strains are highlighted in red. Created in BioRender. Schubert, C. (2026) https://BioRender.com/3p40qc2. **c-e,** Fecal colonization levels of *S.* Typhimurium wild type and mutants deficient in sugar acid utilization pathways following oral infection. Bacterial loads (CFU per gram feces) were determined for wild type and D-gluconate (Δ*idnK* Δ*gntK*, **c**), D-glucuronate (Δ*uxuB*, **d**), or D-glucarate (Δ*gudD*, **e**) mutants at the indicated days post infection. The inoculum was normalized to CFU per gram. Each dot represents an individual mouse; bars indicate median values. Statistical significance was assessed using the non-parametric Wilcoxon rank-sum test; p-values are indicated (not significant, ns: p ≥ 0.05; *\**: p < 0.05; **\*\*:** p < 0.01).

**Figure 2.**
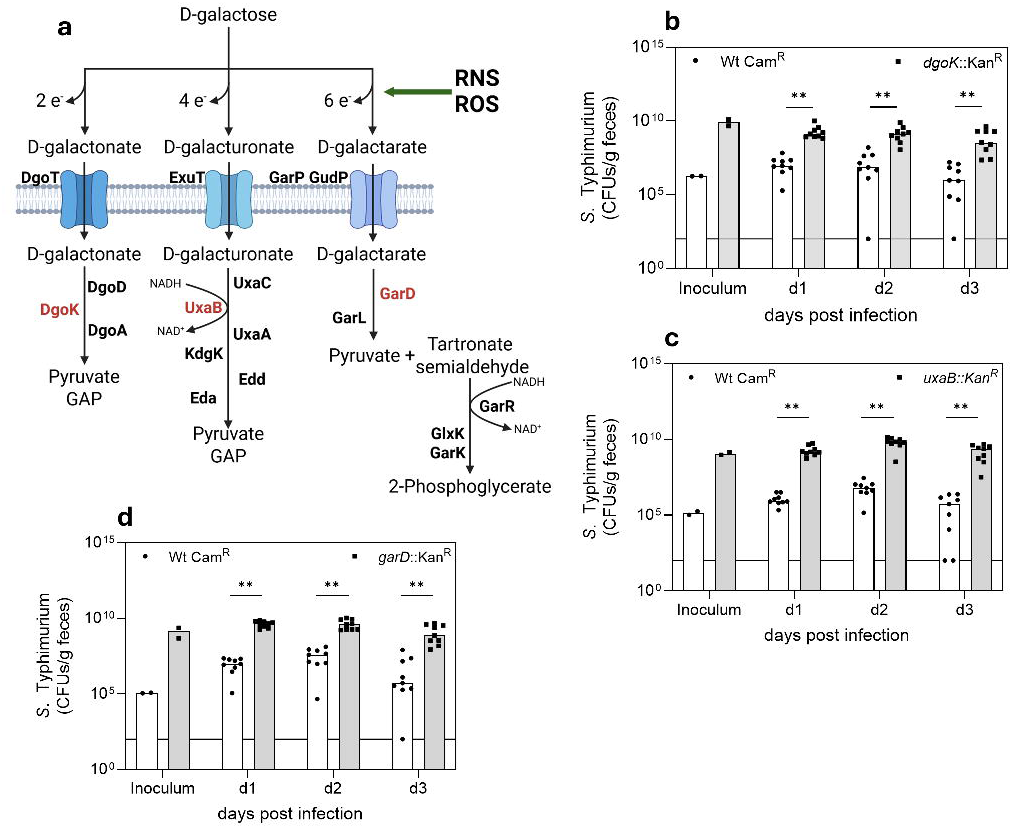
Less contribution of D-galactose-derived sugar acid pathways to wild-type expansion. **a,** Schematic overview of D-galactose-derived sugar acid utilization pathways, including D-galactonate, D-galacturonate, and D-galactarate, and their proposed generation under reactive oxygen and nitrogen species (ROS/RNS). Transporters and metabolic enzymes are indicated; genes targeted in mutant strains are highlighted in red. Created in BioRender. Schubert, C. (2026) https://BioRender.com/3p40qc2. **b–d,** Fecal colonization levels of *S.* Typhimurium wild type (Cam^R^) and mutants deficient in D-galactonate (Δ*dgoK*, **b**), D-galacturonate (Δ*uxaB*, **c**), or D-galactarate (Δ*garD*, **d**) utilization following oral infection of streptomycin-pretreated C57BL/6J mice. Bacterial loads are shown as CFU per gram feces at the indicated days post infection. The inoculum was normalized to CFU per gram. Each dot represents an individual mouse; bars indicate median values. Statistical significance was assessed using the non-parametric Wilcoxon rank-sum test; p-values are indicated (not significant, ns: p ≥ 0.05; *\**: p < 0.05; **\*\*:** p < 0.01).

## Results

### D-gluconate is a key niche promoting *S*. Typhimurium expansion

Classically, competition experiments assess mutant fitness using 1:1 infection mixtures of wild type and mutant strains. To instead evaluate whether a specific sugar acid constitutes an exploitable niche for *S*. Typhimurium wild type, we employed an alternative approach: the mutant strain is introduced in large excess, while the wild type comprises only 0.1% of the inoculum. This setup allows us to test whether the wild type can capitalize on the niche left unoccupied by the mutant and expand to comparable levels over time - as previously demonstrated for D-galactitol utilization ^25^, and several other monosaccharides^18^. By shifting the focus from mutant fitness to wild type expansion, this strategy helps avoid confounding pleiotropic effects often associated with metabolic mutants ^36,37^. Specific-pathogen-free (SPF) C57BL/6J mice were pretreated with streptomycin prior to oral infection with a mixed inoculum of *S.* Typhimurium wild type and the respective mutant strain (0.1% and 99.9%, respectively; 5x10^7^ CFU in total). Bacterial loads were enumerated for 3 days post-infection (**Fig. 1a**). First, we constructed a set of mutants to assess the degradation pathways of D-glucose-derived sugar acids: D-gluconate, D-glucuronate, and D-glucarate. These compounds primarily differ in their degree and position of oxidation - at the C1 position (2 electrons, aldonic acid, D-gluconate), the C6 position (4 electrons, uronic acid, D-glucuronate), or both C1 and C6 (6 electrons, aldaric acid, D-glucarate), respectively. We selected *idnK* and *gntK* (encoding a kinase), *uxuB* (a reductase), and *gudD* (a dehydratase) to test the respective D-glucose-derived sugar acid catabolic pathways (**Fig. 1b**). Two kinases - IdnK and GntK - are capable of phosphorylating D-gluconate in the penultimate step before entry into the Entner–Doudoroff pathway.

We hypothesized that wild-type *S.* Typhimurium fails to reach the stool densities of the respective Δ*idnK* or Δ*gntK* single mutant due to functional redundancy between the two kinases (**Supplementary Fig. 1a, b**). Therefore, we used a Δ*idnK gntK*::Kanᴿ double mutant to block this step entirely. Unlike the single Δ*idnK* or Δ*gntK* mutants, wild-type *S*. Typhimurium successfully catches up to the double Δ*idnK* Δ*gntK* mutant, indicating redundancy in D-gluconate phosphorylation and confirming this metabolic niche as exploitable by *Salmonella* (**Fig. 1c**). Wild-type *S*. Typhimurium was unable to reach comparable levels when competing with the Δ*uxuB* (D-glucuronate) and Δ*gudD* (D-glucarate) mutants, in contrast to the Δ*idnK gntK*::Kanᴿ double mutant (**Fig. 1d, e**). This suggests that D-gluconate represents the most favorable metabolic niche among D-glucose-derived sugar acids in the streptomycin-pretreated mouse model.

### D-galactose–derived sugar acid forms do not support wild-type *S*. Typhimurium expansion

We continued to assess the degradation pathways of D-galactose-derived sugar acids: D-galactonate, D-galacturonate, and D-galactarate. As previously explained for D-glucose, these compounds primarily differ in their degree and position of oxidation - at the C1 position (D-galactonate), the C6 position (D-galacturonate), or both C1 and C6 (D-galactarate), respectively. We selected *dgoK* (encoding a kinase), *uxaB* (a reductase), and *garD* (a dehydratase) (**Fig. 2a**). We again used the streptomycin-pretreated C57BL/6J model to test whether wild-type *S*. Typhimurium can exploit the niche created by the respective mutant to ecologically invade the mutant population (**Fig. 1a**). In contrast to D-glucose-derived sugar acids, D-galactose-derived counterparts did not support wild-type *S*. Typhimurium expansion to stool densities comparable to the respective mutant strains (**Fig. 2b-d**). Interestingly, only for D-galacturonate (Δ*uxaB*) and D-galactarate (Δ*garD*), the wild type showed a trend toward increased abundance between day 1 and day 2 post infection, but numbers collapsed by day 3 (**Fig. 2c, d**). In contrast to D-glucose-derived sugar acids, particularly D-gluconate, D-galactose-derived sugar acids did not fully support ecological expansion of wild-type *S*. Typhimurium into the mutant population.

We also tested *uxaC* and *kdgK*, encoding an isomerase and a kinase, which are shared enzymes in the degradation of D-glucuronate and D-galacturonate (**Fig. 1b, and Fig. 2a**). Only in the case of the *uxaC* mutant was wild-type *S*. Typhimurium able to partially expand to mutant levels, likely reflecting the greater exploitability of D-glucuronate compared with D-galacturonate (**Supplementary Fig. 2**). In conclusion, D-glucose-derived sugar acids, in particular D-gluconate, provide a metabolic niche sufficient to promote *S.* Typhimurium expansion. In contrast, D-galactose-derived sugar acids do not constitute an equally effective exploitable niche for *S*. Typhimurium in streptomycin-pretreated mice.

### D-glucose–derived sugar acids are the key driver of wild-type *S*. Typhimurium expansion

We followed up on these findings by generating *S*. Typhimurium multimutants targeting degradation pathways for all three D-glucose- and D-galactose-derived sugar acids. The Δ*idnK* Δ*gntK* Δ*gudD uxuB*::Kanᴿ (abbreviated: Δ4 D-glc) strain lacks the ability to utilize D-gluconate, D-glucarate, and D-glucuronate. Surprisingly, in our ecological invasion experiment the wild-type strain expanded to comparable stool densities as the Δ4 D-glc mutant as early as day 1 post-infection (**Fig. 3a**). Previous studies have shown that multimutants can facilitate faster expansion of wild-type *Salmonella* due to niche availability but may also be prone to displacement if the fitness cost of the mutations is too severe^18^. The latter was not observed here; the Δ4 D-glc multimutant remained at a stable abundance throughout the first two days and declined only slightly by day 3 (**Fig. 3a**). In addition, we assessed the triple mutant lacking both D-gluconate and D-glucarate utilization (Δ*idnK* Δ*gntK gudD*::Kanᴿ) and observed an intermediate phenotype between the D-gluconate mutant (Δ*idnK* Δ*gntK*) and the Δ4 D-Glc mutant for wild type expansion (**Supplementary Fig. 3a**). The Δ*dgoK* Δ*uxaB garD*::Kanᴿ (alternative name: Δ3 D-gal) strain is deficient in the utilization of D-galactonate, D-galacturonate, and D-galactarate. In contrast to Δ4 D-glc, wild-type *S.* Typhimurium was could not efficiently expand when competing against this mutant (**Fig. 3b**). This suggests that D-galactose-derived sugar acids fail to support robust expansion, whereas their D-glucose counterparts provide an exploitable niche for wild-type *S*. Typhimurium in the streptomycin-pretreated mouse model. This suggests a strain-dependent phenotype, as previous work showed that a mutant defective in D-galactarate utilization (Δ*garD*) exhibit a severe fitness defect at 4 days post infection in ATCC14028s, another frequently used laboratory strain of *S*. Typhimurium ^29^. This suggests that different *S*. Typhimurium strains exhibit distinct sugar acid preferences. Considering the previous results, D-gluconate plays the dominant role among D-glucose-derived sugar acids in providing *Salmonella* SL1344 with an exploitable niche that allows expansion to mutant levels.

**Figure 3.**
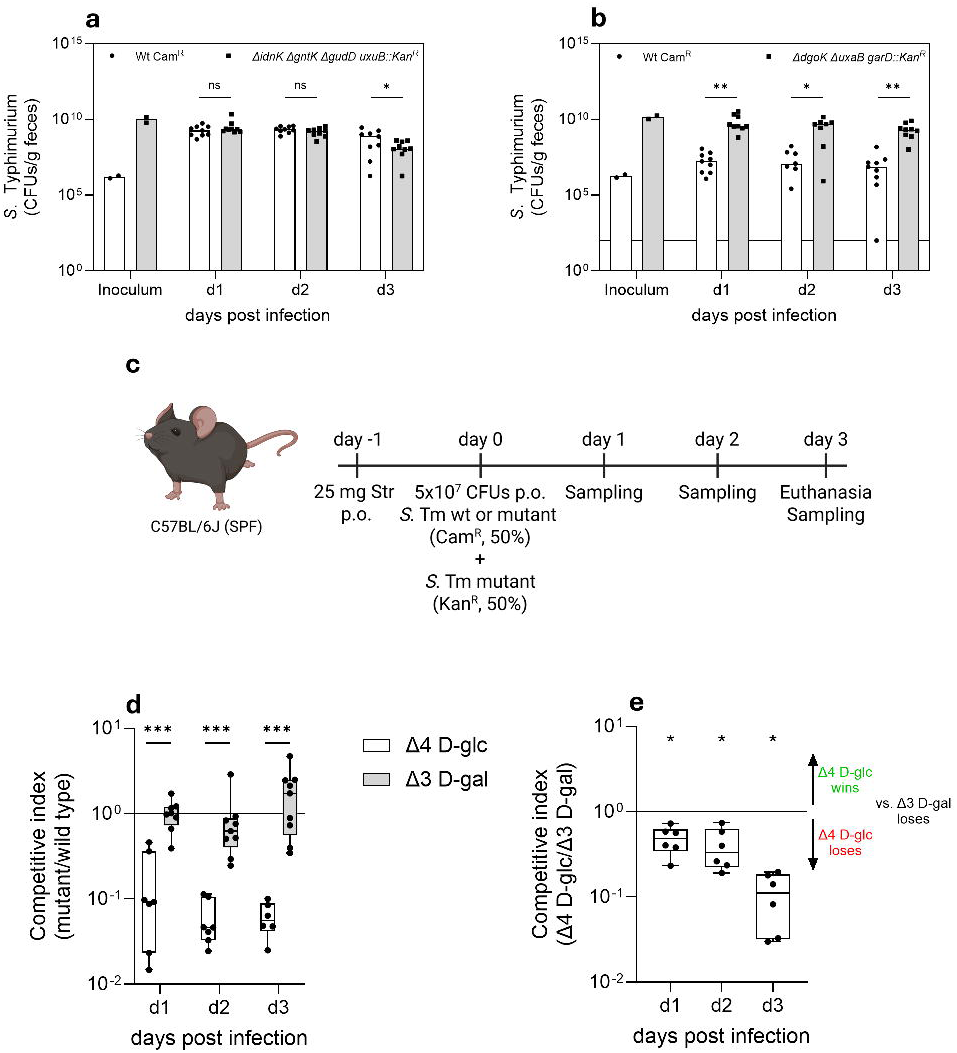
D-glucose, but not D-galactose-derived sugar acids, are an exploitable niche to *S*. Typhimurium during gut colonization. **a,b**, Fecal colonization levels of *S*. Typhimurium wild type (Cam^R^) and mutants lacking combined D-glucose-derived (Δ*idnK* Δ*gntK* Δ*gudD* Δ*uxaB*, Δ4 D-glc, **a**) or D-galactose-derived (Δ*dgoK* Δ*uxaB* Δ*garD*, Δ3 D-gal, **b**) sugar acid utilization pathways following oral infection of streptomycin-pretreated C57BL/6J mice. Bacterial loads are shown as CFU per gram feces at the indicated days post infection. The inoculum was normalized to CFU per gram. **c**, Experimental design of the competitive infection assay. Streptomycin-pretreated C57BL/6J mice were orally inoculated with a 1:1 mixture of *S*. Typhimurium wild type (Cam^R^) and mutant strains (Kan^R^). Fecal samples were collected daily, and animals were euthanized on day 3 post infection. Created in BioRender. Schubert, C. (2026) https://BioRender.com/3p40qc2. **d**, Competitive indices of mutants lacking D-glucose-derived (Δ4 D-glc, white) or D-galactose-derived (Δ3 D-gal, grey) sugar acid utilization pathways relative to wild type over the course of infection. **e**, Competitive indices of Δ4 D-glc relative Δ3 D-gal, indicating a dominant contribution of D-glucose-derived sugar acid utilization to competitive fitness. Each dot represents an individual mouse; bars or box plots indicate median values. Statistical significance was assessed using the non-parametric Wilcoxon rank-sum test for subpanels **a**, **b**, and **e**, and a two-tailed Mann–Whitney U test for subpanel **d**; p-values are indicated (not significant, ns: p ≥ 0.05; *\**: p < 0.05; **\*\*:** p < 0.01; **\*\*\*:** p < 0.001)

We next assessed the fitness of Δ4 D-glc and Δ3 D-gal mutants in classical 1:1 competition with wild-type *S.* Typhimurium in the streptomycin-pretreated C57BL/6J model, shifting the focus from previous ecological invasion experiments to mutant fitness during gut-luminal colonization (**Fig. 3c**). While Δ3 D-gal showed no fitness loss throughout the experiment, Δ4 D-glc dropped tenfold by day 1 post-infection and remained at this level, thereafter, indicating a modest fitness attenuation during early gut colonization (**Fig. 3d**). This trend was also evident when comparing absolute bacterial loads: Δ4 D-glc remained consistently lower than wild-type *S*. Typhimurium, whereas Δ3 D-gal did not differ from the wild type (**Supplementary Fig. 3b, c**). Even in the presence of an additional niche competitor, *E. coli* 8178, previously shown to compete with *S*. Typhimurium for shared nutrient resources ^22,23^, the Δ4 D-glc mutant’s fitness remained unaffected (**Supplementary Fig. 3d**).

To determine which metabolic niche is more important for *S*. Typhimurium, we introduced neutral genetic barcodes into the Δ4 D-glc and Δ3 D-gal mutants, each carrying either kanamycin or chloramphenicol resistance markers chromosomally, enabling direct comparison in a classical 1:1 competitive infection experiment. Akin to the competition with wild-type *S.* Typhimurium, Δ4 D-glc displayed significantly reduced fitness compared with Δ3 D-gal, with the difference becoming more pronounced by day 3 post-infection (**Fig. 3e**). This trend was consistent when comparing the absolute loads of Δ4 D-glc and Δ3 D-gal (**Supplementary Fig. 3e**). D-glucose-derived sugar acids provide an exploitable niche for *Salmonella*; however, the inability to utilize this niche surprisingly confers a modest fitness cost. To follow up on this observation, we infected streptomycin-pretreated C57BL/6J mice in a single-infection experiment with either the Δ4 D-glc mutant or wild-type *S*. Typhimurium. The only significant difference was observed on day 1 post-infection and was no longer detectable at later time points, indicating no sustained fitness defect in single-infection experiments (**Supplementary Fig. 4a**). To evaluate inflammation, fecal lipocalin-2 levels were monitored daily post-infection. Lipocalin-2 is a sensitive marker of gut inflammation, and no significant differences were observed between wild-type and Δ4 D- glc infections, indicating comparable inflammatory responses (**Supplementary Fig. 4b**). This was further confirmed by histopathological analysis of cecal tissue on day 3 post-infection (**Supplementary Fig. 4c**). Although the transient difference observed on day 1 post-infection underscores a role for D-glucose-derived sugar acids during early gut colonization, sugar acids overall represent an exploitable but nonessential niche for *Salmonella*, even in competitive settings in short-term infection models.

### Long-term infection severely impairs D-glucose-derived sugar mutants

Previous experiments in C57BL/6J mice revealed a significant 10-fold fitness defect of the Δ4 D-glc mutant during early gut colonization (**Fig. 3d**). We therefore focused on this mutant to assess the role of D-glcuose-derived sugar acids during long-term infection in the 129S6/SvEvTac mouse model. A key distinction between these models is the Nramp1 (Slc11a1) genotype. C57BL/6J mice carry a non-functional allele, rendering them highly susceptible to systemic *S*. Typhimurium infection, whereas 129S6/SvEvTac mice harbor a functional Nramp1 allele that confers increased resistance to systemic dissemination ^38^. An advantage of this model is that infection can be studied for longer than 4 days, allowing analysis of how *S.* Typhimurium colonization unfolds beyond the onset of acute *Salmonella*-induced inflammation ^39^. Similarly, 129S6/SvEvTac mice were pretreated with streptomycin and subsequently inoculated with *S*. Typhimurium. Fecal shedding were monitored over the 9-day infection period, including systemic loads at the end of the infection (**Fig. 4a**). We first assessed the competitive behavior of the Δ4 D-glc mutant, which is deficient in the utilization of all D-glucose-derived sugar acids, in classical 1:1 competition experiments with the wild type. During the first 3 days of infection, we observed a phenotype similar to that in the C57BL/6J model, with mutant fitness reduced by approximately 10- to 50-fold relative to the wild type. This attenuation plateaued at around 100-fold by day 7 post-infection and subsequently declined further, ultimately reaching a 1,000-fold reduction compared with the wild type (**Fig. 4b**). The streptomycin-induced effects on iNOS activity and sugar acid production likely subside within the first 3–4 days post treatment. The continued decline in the competitive index beyond day 3 therefore suggests that intestinal inflammation itself sustains sugar acid–dependent growth. Notably, this severe attenuation was not observed in systemic dissemination, where the Δ4 D-glc mutant exhibited only an approximately 5- to 10-fold fitness defect in the spleen, liver, and mesenteric lymph nodes (**Fig. 4c**). In conclusion, D-glucose-derived sugar acid metabolism is more critical for gut luminal growth than for systemic dissemination. When the wild type was excluded and infections were performed with the Δ4 D-glc mutant alone, we observed no fitness defect relative to the wild type (**Fig. 4d**). In the absence of wild-type *S*. Typhimurium, the Δ4 D-glc mutant colonized stably throughout the 9-day infection period. The same observations were made for the systemic sites, spleen, liver, and mesenteric lymph nodes (**Fig. 4e**). This finding aligns with our previous observations showing that the Δ4 D-glc mutant maintains wild-type fitness in single-infection experiments. However, once a niche competitor is present, such as wild-type *S*. Typhimurium, D-glucose-derived sugar acids become increasingly important for sustained colonization during inflammation in a long-term infection model.

**Figure 4.**
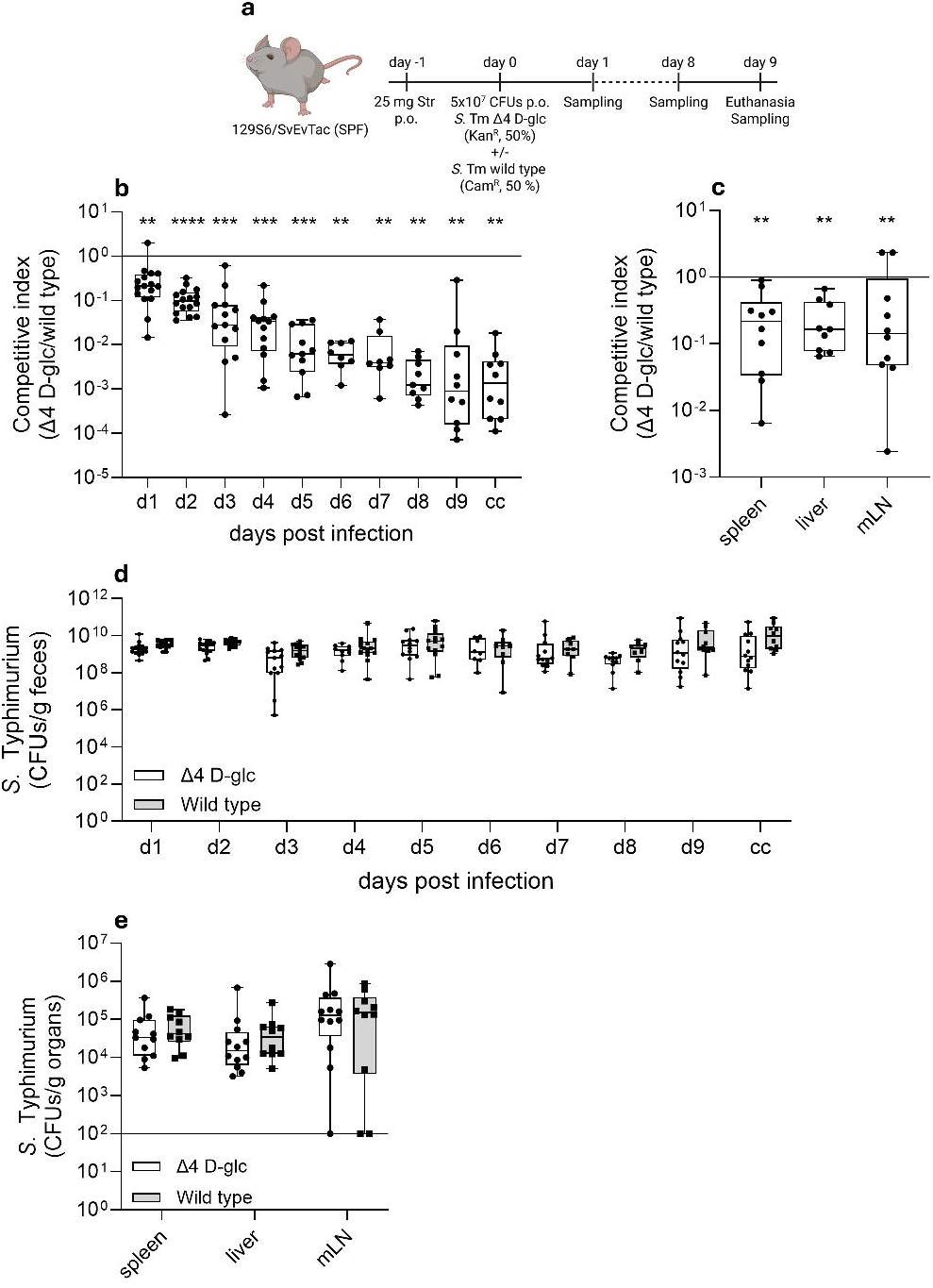
Attenuated fitness of the Δ4 D-glc mutant during long-term infection in 129S6/SvEvTac mice. (**a**) Experimental design. Specific pathogen-free 129S6/SvEvTac mice were pretreated with streptomycin and subsequently inoculated with a 5×10⁷ CFU mixture of *S.* Typhimurium wild type and the Δ4 D-glc mutant, or with the Δ4 D-glc mutant alone. Fecal samples were collected at the indicated time points, and mice were euthanized on day 9 post infection for analysis of systemic dissemination. Created in BioRender. Schubert, C. (2026) https://BioRender.com/3p40qc2. **b,** Competitive index of the Δ4 D-glc mutant relative to the wild type in feces and cecum content at the indicated days post infection. A value of 1 indicates equal fitness. **c**, Competitive index of the Δ4 D-glc mutant relative to the wild type in systemic sites - spleen, liver, and mesenteric lymph nodes (mLN) - at day 9 post infection. **d**, Fecal bacterial loads of *S.* Typhimurium over the course of infection. Each data point represents one mouse; box plots show median and interquartile range. **e**, Absolute bacterial loads of *S.* Typhimurium in systemic organs at day 9 post infection. Each symbol represents an individual mouse. Statistical significance was assessed using the non-parametric Wilcoxon rank-sum test; p-values are indicated (not significant, ns: p ≥ 0.05; *\**: p < 0.05; **\*\*:** p < 0.01; **\*\*\*:** p < 0.001; **\*\*\*\*:** p < 0.0001).

### LPS-induced oxidative stress does not impair sugar acid utilization by *S*. Typhimurium

Streptomycin pretreatment induces elevated expression of iNOS in the cecal mucosa, leading to the production of reactive nitrogen species (RNS) and subsequent oxidation of monosaccharides ^29^. Recently, a mouse model was characterized in which lipopolysaccharides (LPS) derived from *S*. Typhimurium activate Toll-like receptor 4 (TLR4), triggering a TLR4-dependent increase in gut-luminal reactive oxygen species (ROS). Both treatments promote *Enterobacteriaceae* expansion, although LPS-induced colonization results in bacterial loads approximately 100-fold lower at the peak compared to streptomycin pretreatment ^40^. However, the electron donors fueling these blooms have remained unknown. Based on the observations above, we hypothesized that LPS might trigger the production of sugar acids that fuel gut-luminal *Enterobacteriaceae* blooms.

In the LPS injection model, specific-pathogen free C57BL/6J mice are administered LPS either intravenously (i.v.) or intraperitoneally (i.p.) concurrently with *S.* Typhimurium inoculation. Mice were euthanized the following day, and bacterial loads were quantified on selective MacConkey agar plates (**Supplementary Fig. 5a**). LPS was originally administered intravenously (i.v.) (**Supplementary Fig. 5b**) ^40^. We adapted this model by delivering LPS intraperitoneally (i.p.), which resulted in a comparable *S.* Typhimurium bloom relative to the PBS control (**Supplementary Fig. 5c**). We next tested mutants defective in the degradation of D-glucose- and D-galactose-derived sugar acids in 1:1 competition experiments against wild-type *S.* Typhimurium using the LPS mouse model (**Supplementary Fig. 5d,e**). We opted for 1:1 competition assays instead of ecological invasion assays due to the overall lower stool densities. In both cases, no significant fitness defect was observed compared to the wild type. We then proceeded with multimutants lacking the full set of degradation pathways for either D-glucose-derived sugar acids (Δ4 D-glc) and D-galactose-derived sugar acids (Δ3 D-gal). Similar to the single mutants, both strains exhibited wild-type-like fitness (**Supplementary Fig. 5f**). Interestingly, as observed previously in the streptomycin-pretreated mouse model, the Δ4 D-glc mutant exhibited a slight but significant fitness disadvantage relative to the Δ3 D-gal mutant in a competition experiment (**Supplementary Fig. 5g**). In conclusion, LPS, which is known to trigger ROS production, does not impair the fitness of *S*. Typhimurium sugar acid mutants. Notably, D-glucose-derived sugar acids appear to be slightly more important than the D-galactose counterparts, as reflected by the modest fitness defect of the Δ4 D-glc mutant compared with the Δ3 D-gal mutant (**Supplementary Fig. 5g**).

### D-gluconate supported the highest *in vitro* growth of *S*. Typhimurium

We demonstrate that D-glucose-derived sugar acids represent an exploitable niche that promotes *S.* Typhimurium expansion in the streptomycin-pretreated C57BL/6J model, with D-gluconate identified as the key sugar acid driving *Salmonella* colonization. The inability of D-galactose-derived sugar acids to support *S*. Typhimurium colonization may be due to two factors: either they are poor carbon sources, or their concentrations in the gut are insufficient. To address the first possibility, we cultured wild-type *S.* Typhimurium *in vitro* using a selection of commercially available D-glucose- and D-galactose-derived sugar acids as sole carbon sources in minimal medium under aerobic conditions (**Fig. 5a, b**). D-gluconic acid readily forms a stable δ-lactone ring that equilibrates with the open-chain form in water. In contrast, D-galactonic acid produces multiple, less stable lactone isomers that interconvert and hydrolyze unpredictably. Moreover, calcium galactonate - the most stable commercial form - has very poor water solubility, making it unsuitable for defined growth assays. For these reasons, we limited our analyses to D-galacturonate and D-galactarate among the D-galactose-derived sugar acids. *S*. Typhimurium was grown overnight in minimal medium supplemented with pyruvate as a neutral carbon source. The following day, subcultures were inoculated into fresh minimal medium containing the indicated sugar or sugar acid as the sole carbon source. *S*. Typhimurium was able to grow on D-glucose, D-gluconate, and D-glucuronate, and to a lesser extent on D-glucarate (**Fig 5a**). However, a distinct shift in lag phase was observed: growth initiated earlier on D-glucose compared to D-gluconate and D-glucuronate (**Supplementary Fig. 6a**). In contrast, D-galactose-derived sugar acids failed to support *S*. Typhimurium growth, unlike D-galactose itself, which did (**Fig. 5b, Supplementary Fig. 6b**).

**Figure 5.**
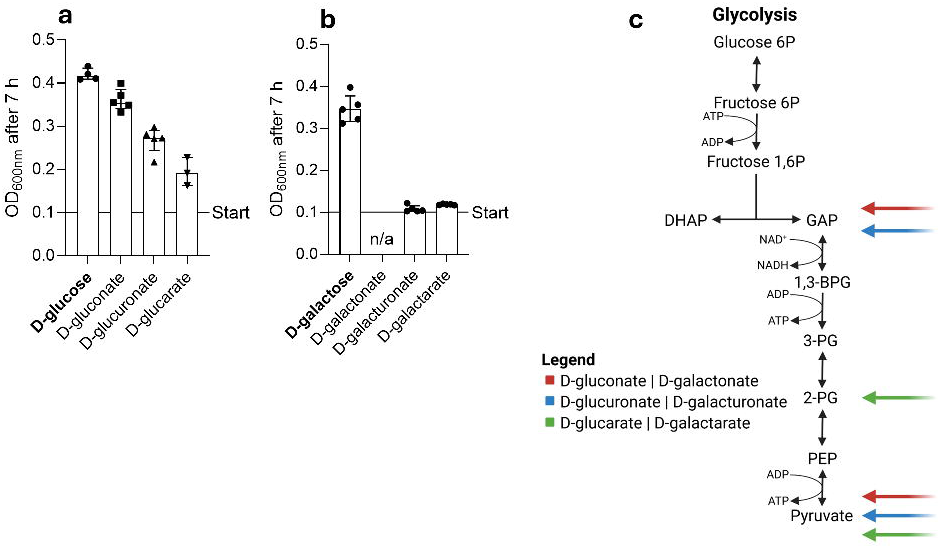
Differential growth on sugar acids and integration into central carbon metabolism. a,b, Growth of *S.* Typhimurium in minimal medium supplemented with the indicated sugar acids derived from D-glucose (**a**) or D-galactose (**b**). Optical density (OD_600_) is plotted for 7 h of incubation. Start indicates the initial OD_600_ at inoculation; n/a, D-galactonate was not available. Data are shown as median **c,** Schematic representation of the entry points of D-gluconate, D-glucuronate, and D-glucarate catabolism, as well as D-galactose-derived sugar acids, into central carbon metabolism. Colored arrows indicate the metabolic nodes at which each sugar acid feeds into glycolysis or lower glycolytic intermediates, highlighting differences in pathway integration. Created in BioRender. Schubert, C. (2026) https://BioRender.com/3p40qc2.

In conclusion, while D-glucose-derived sugar acids can support growth of *S*. Typhimurium SL1344 as sole carbon sources, this is not the case for D-galactose-derived sugar acids. Interestingly, D-galactarate specifically was shown to promote growth of the ATCC14028s wild type ^29^, again indicating strain-dependent sugar acid utilization patterns. D-gluconate, D-glucuronate, and D-glucarate are progressively more oxidized forms of D-glucose. Because much of the oxidative potential that normally drives ATP production is already expended, these substrates enter central metabolism - primarily glycolysis - at downstream entry points, resulting in a proportionally lower ATP yield (**Fig. 5c**) ^30^. In principle, both D-gluconate and D-glucuronate are metabolized to pyruvate and glyceraldehyde-3-phosphate (GAP). In contrast, D-glucuronate must first be reduced, consuming NADH and thereby yielding less net energy under aerobic conditions (**Fig. 1b**). This is also true for D-galactose-derived sugar acids; however, D-galacturonate and D-galactarate fail to support growth comparable to their D-glucose-derived sugar acids counterparts. Although the degradation pathways of D-glucuronate and D-galacturonate largely converge, differences in enzyme activity and substrate uptake likely result in distinct metabolic efficiencies. These differences appear to be sufficient to generate measurable growth disparities under nutrient-limited conditions, as observed in this study.

### D-gluconate is the most abundant sugar acid

D-glucose-derived sugar acids, particularly D-gluconate, constitute the dominant sugar acid substrates supporting *S*. Typhimurium expansion in the streptomycin-pretreated mouse model, with an especially pronounced role during long-term infection. This finding implies that these substrates are readily available in gut-luminal contents. To test this, we collected cecal contents from untreated specific pathogen–free (SPF), streptomycin-pretreated, and LPS-treated C57BL/6J mice and analyzed monosaccharide levels by LC–MS (**Fig. 6a**). As previously reported ^11,18^, D-glucose and D-galactose levels are generally in the low millimolar range. D-glucose- and D-galactose-derived sugar acid concentrations were generally highest in the streptomycin-pretreated model, with the corresponding aldonic acids, D-gluconate and D-galactonate, showing the highest levels (**Fig. 6a**). Notably, D-gluconate concentrations even exceeded those of D-glucose. Chemically, aldonic acids represent the lowest-energy oxidation products of aldoses, requiring only a two-electron oxidation at C1. Under conditions of oxidative stress, such partial oxidation reactions are favored, providing a plausible explanation for the preferential accumulation of aldonic acids such as D-gluconate and D-galactonate in the gut lumen. The lack of sugar acid accumulation in the LPS model may reflect a predominance of ROS-driven inflammation ^40^, which is less effective at promoting stable aldose oxidation in the gut lumen than the RNS-mediated oxidation of the streptomycin-pretreated model, which depends on the NOS2 pathway ^29^. In addition to the sugar acid forms of D-glucose and D-galactose, we analyzed additional monosaccharides (**Supplementary Fig. 7a**). As previously reported ^18^, streptomycin treatment also increased the pentoses L-arabinose and D-xylose, a trend that was recapitulated in this dataset. In contrast, LPS-treated mice exhibited a monosaccharide profile largely similar to that of untreated SPF mice. While the two sugar acids L-idonate and D-mannuronate were also increased in cecal contents in streptomycin-pretreated mice, their origin is likely similar to that of the major D-glucose-derived sugar acids (**Supplementary Fig. 7a**). L-idonate derives from oxidation of the rare hexose L-idose or from epimerization of D-glucuronate, whereas D-mannuronate represents the uronic acid of D-mannose, formed by oxidation at the C6 position. The comparatively low concentration of these substrates is consistent with the limited availability of their precursor hexoses in the gut lumen, particularly D-mannose. In conclusion, D-glucose is not only the most abundant dietary monosaccharide, but it is also prevalent in the cecal contents of mice. As D-gluconate represents the chemically simplest oxidation product of D-glucose, we detected it at millimolar concentrations in the gut lumen (**Fig. 6a**). However, this was observed only in the streptomycin-pretreated model, as LPS-induced oxidative stress alone appears insufficient to drive substantial sugar acid formation in the gut lumen.

**Figure 6.**
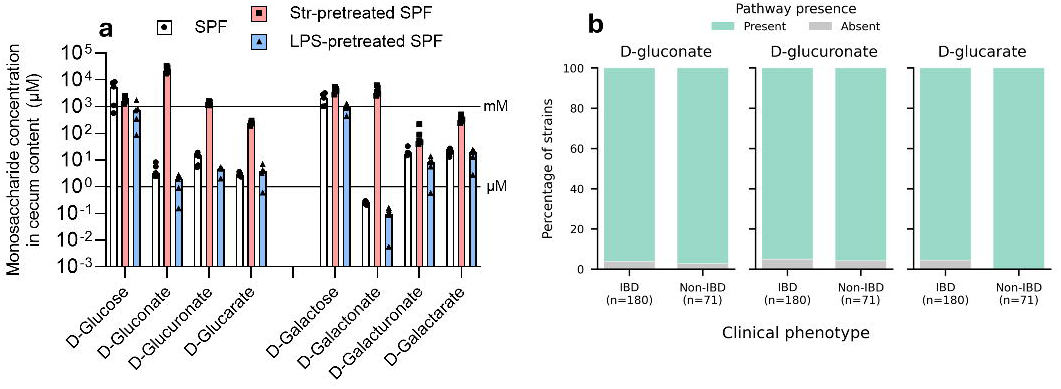
Sugar acids are present in the murine cecum at physiologically relevant concentrations under different inflammatory conditions. **a,** Concentrations of monosaccharides and corresponding sugar acids in cecal contents of specific-pathogen-free (SPF), streptomycin-pretreated SPF, and LPS-pretreated SPF mice. Shown are D-glucose and D-galactose together with their derived sugar acids D-gluconate, D-glucuronate, D-glucarate, D-galactonate, D-galacturonate, and D-galactarate. Data are displayed on a logarithmic scale; horizontal reference lines indicate micromolar (µM) and millimolar (mM) concentration ranges. Each symbol represents an individual mouse; bars indicate median values. **b**, Bar plots show the percentage of *E. coli* strains harboring the indicated metabolic pathways in isolates derived from patients with inflammatory bowel disease (IBD, n = 180) and non-IBD controls (n = 71). Pathway presence was determined based on genomic analysis. For all three pathways, the vast majority of strains in both cohorts encode the complete degradation pathways, with only a small fraction lacking the respective genes. No substantial differences in pathway prevalence were observed between IBD and non-IBD-derived isolates.

### Genes involved in sugar acid utilization are prevalent in both healthy and IBD states

Inflammatory bowel disease (IBD) is a chronic relapsing inflammatory disorder of the gastrointestinal tract, primarily encompassing Crohn’s disease and ulcerative colitis. Its pathogenesis arises from a dysregulated immune response to the intestinal microbiota in genetically susceptible hosts ^41,42^. Inflammation profoundly alters gut physiology and nutrient availability, reshaping microbial community structure and function. A key driver is inducible nitric oxide synthase (NOS2), which is strongly upregulated via NF-κB signaling ^43^ and elevated 4 to 12-fold in patients with active IBD ^31–33^, suggesting oxidation of gut monosaccharides. Therefore, we investigated sugar acid utilization in metagenomic datasets from individuals with IBD and non-IBD controls. We focused on metabolic pathways prevalent specifically in *Escherichia coli* and quantified the abundance of these pathways across metagenomic assembled and isolate genomes (**Fig. 6b**). We observed no difference in the presence of D-glucose- or D-galactose-derived sugar acid metabolic pathways in *E. coli* between IBD patients and non-IBD controls based on genomic analysis (**Fig. 6b** and **Supplementary Fig. 7b**). Low-abundance *E. coli* are almost universally present in healthy individuals ^44^. Certain *E. coli* lineages are classic examples of pathobionts: they persist at low abundance in healthy individuals but can expand and contribute to pathology in inflammatory bowel disease ^45^. Importantly, inflammation selects for pre-existing metabolic pathways that confer a fitness advantage and drive bacterial expansion under these conditions, as previously demonstrated for the utilization of inorganic electron acceptors such as oxygen ^14^, nitrate ^13^, and tetrathionate ^12^. This clearly establishes D-glucose-derived sugar acids, particularly D-gluconate, as an important inflammation-dependent nutrient niche for pathogens and pathobionts.

## Discussion

In this study, we demonstrate that D-gluconate, and more broadly D-glucose-derived sugar acids including D-gluconate, D-glucuronate, and D-glucarate, constitute an exploitable nutrient niche for *S.* Typhimurium during early gut colonization of streptomycin-pretreated mice (**Fig. 1**, **Fig. 3**) and also contribute to long-term infection in the inflamed intestine (**Fig. 4**). In line with D-glucose being the most abundant dietary monosaccharide, LC-MS analysis of cecal samples revealed that D-gluconate is produced in high amounts in the gut of streptomycin-pretreated mice (**Fig. 6a**). In contrast, D-galactose-derived sugar acids contribute little to gut-luminal *S*. Typhimurium growth, consistent with the poor *in vitro* growth observed for different D-galactose-derived sugar acids when used as the sole carbon source (**Fig. 5b**). In a mouse model characterized by milder ROS-mediated oxidative stress ^40^, sugar acid (multi)mutants showed no fitness defect (**Supplementary Fig. 5**), confirming that the level of oxidative stress elicited by a sub-lethal dose of LPS is insufficient to drive substantial monosaccharide oxidation (**Fig. 6a**). It is tempting to speculate that monosaccharide oxidation may be particularly relevant in chronically inflamed conditions such as inflammatory bowel disease (IBD). Bioinformatic analysis of metagenomic data revealed that sugar acid metabolism genes are equally prevalent in samples from IBD patients and healthy individuals (**Fig. 6b**), underscoring the pathobiont nature of *E. coli*, which maintains a metabolic repertoire that enables proliferation when ecological niches open within the gut, similar to its capacity for microaerophilic and anaerobic respiration ^13,46–49^.

Monosaccharides are abundant in the diet and are therefore present at substantial levels across a wide range of mouse models that differ in microbiome complexity ^11,16,18,50^. Sugar acids are relatively scarce under homeostatic conditions ^51^ but become abundant during inflammation or antibiotic pretreatment (**Fig. 6a**). A central driver of their formation is inducible nitric oxide synthase, encoded by NOS2, which is strongly upregulated via NF-κB signaling during inflammatory responses ^43^ and is elevated 4 to 12-fold in patients with active IBD ^31–33^. iNOS-derived nitric oxide promotes oxidative conversion of D-glucose and D-galactose into sugar acids such as D-glucarate and D-galactarate, creating a nutrient landscape that selects for *Enterobacteriaceae* ^52–55^ and drives convergent evolution of D-glucarate and D-galactarate metabolism in both commensal and pathogenic microbes ^34^. Importantly, antibiotic pretreatment in the streptomycin mouse model also induces iNOS expression and oxidative stress ^29^. Thus, the streptomycin model recapitulates a key mechanistic feature of the IBD gut environment, namely iNOS-dependent sugar oxidation, making it a representative system to study inflammation-associated metabolic niches. However, in this study we shift the focus to D-gluconate, a more favorable nutrient source that is likely present at higher concentrations in IBD and therefore represents a particularly relevant substrate for *Enterobacteriaceae*, as its degradation pathway is conserved in the core genome of this family ^35^. The relevance of D-gluconate for *K. pneumonia*e was recently demonstrated ^5^. Together with our new data, this confirms that this sugar acid is a common carbohydrate source fueling *Enterobacteriaceae* growth in host-associated environments.

D-glucose and D-galactose can stably generate all three classes of sugar acids, namely aldonic, uronic, and aldaric acids. Among these, aldonic acids are energetically the most favorable because they are only partially oxidized and therefore retain the highest ATP-generating potential (**Fig. 5c**). Since D-glucose is the most abundant dietary hexose in the gut, its C1 oxidation product, D-gluconate, is the predominant sugar acid formed during oxidative stress. Because D-gluconate combines high concentration, low oxidative barrier to formation, and the greatest remaining energetic yield among sugar acids, it should represent the most accessible and metabolically rewarding sugar acid niche for enteric bacteria, specifically *Enterobactericeae*. Consistent with this, D-gluconate provides the strongest growth advantage and likely constitutes the primary sugar acid encountered by *S*. Typhimurium during inflammation. Therefore, controlling the availability or formation of D-gluconate in the gut may offer new opportunities for anti-infective strategies and preventive interventions. Our findings provide a foundation for future research in this direction.

## Acknowledgements

We would like to acknowledge and thank the staff at the ETH animal facilities (EPIC and RCHCI; especially Manuela Graf, Katharina Holzinger, Dennis Mollenhauer, Sven Nowok, Samuel Boateng, and Dominik Bacovcin), and extend many thanks to members of the Hardt lab, as well as the NCCR Microbiomes, for their helpful comments and discussions. In addition, we would like to thank Lena Ernstberger, Davide Mazzali, and Vincent Peuker for their practical contributions. This work has been funded by grants from the Swiss National Science Foundation (310030_192567, 10.001.588 and NCCR Microbiomes grant 51NF40_180575) to W.-D.H.. This work has been further funded by grants from the Swiss National Science Foundation (grant 310030_19256) attributed to N.N. and C.v.M. and NCCR Microbiomes grant 51NF40_180575 to C.v.M..

## Author contributions

C.S. and W.-D.H. conceived and designed the experiments. C.S. performed the *in vivo* experiments with the help of M.H., J.K., S.K. and S.P.. M.H., A.S. and U.S. collected samples and analyzed monosaccharides by LC-MS. N.N. and C.v.M. performed the bioinformatic analysis to identify sugar acid metabolic genes in metagenomic datasets from non-IBD and IBD patients. L.B. performed whole-genome sequencing of the sugar acid multimutants. M.H. and B.D.N. performed histopathological scoring. C.S. wrote the manuscript with contributions from all authors.

## Ethics declaration

### Competing interests

The authors declare no competing interests.

### Data availability

The whole-genome sequencing data, including genomes, are available in the European Nucleotide Archive (ENA) under accession number PRJEB108430. We adhered to the data reuse guidelines presented in ^56^.

## Methods

### Ethics statement

All animal experiments were performed in agreement with the guidelines of the Kantonales Veterinäramt Zürich under licenses ZH108/22, ZH109/2022, and ZH092/2025, and the recommendations of the Federation of European Laboratory Animal Science Association (FELASA).

### Animals

We used male and female mice aged 7-12 weeks, randomly assigning animals of either sex to the experimental groups. The consideration of animal sex was not included in the study design. All mice were maintained on the normal mouse chow (Kliba Nafag, 3537; autoclaved; per weight: 4.5% fat, 18.5% protein, 50% carbohydrates, 4.5% fiber). The mice originated from C57BL/6J and 129S6/SvEvTac breeders initially obtained from Jackson Laboratories. Mice with a normal complex microbiota were specific pathogen-free (SPF) and bred under full barrier conditions in individually ventilated cage systems at the EPIC mouse facility of ETH Zurich (light/dark cycle 12:12 h, room temperature 21±1°C, humidity 50±10%). All studies were conducted in compliance with ethical and legal requirements and were reviewed and approved by the Kantonales Veterinäramt Zürich under licenses ZH108/22, ZH109/2022, and ZH092/2025.

### Bacteria and culture conditions

All *Salmonella* strains are isogenic to *Salmonella* Typhimurium SB300, a re-isolate of SL1344. The names SB300 and SL1344 are used synonymously. They are listed in Supplementary Data 1. All plasmids used in this study are listed in Supplementary Data 2. All strains were routinely grown overnight at 37°C in Lysogeny broth (LB) with agitation. Strains were stored at -80°C in peptone glycerol broth (2% w/v peptone, 5% v/v glycerol (99.7%)). Custom oligonucleotides were synthetized by Microsynth AG (Switzerland) and are listed in Supplementary Data 3.

### Homologous recombination by λ red

Single-gene knockout strains were generated using the λ red single-step protocol ^57^. Primers were designed with an approximately 40 bp overhanging region homologous to the genomic region of interest and 20 bp binding region corresponding to the antibiotic resistance cassette (Supplementary Data 3). PCR amplification was performed using the plasmid pKD4 and pKD13 for kanamycin resistance. DreamTaq Master Mix (Thermo Fisher Scientific) was employed, followed by digestion of the template DNA using FastDigest DpnI (Thermo Fisher Scientific). Subsequently, the PCR product was purified using the Qiagen DNA purification kit (Macherey-Nagel). SB300 with either the pKD46 or pSIM5 plasmid was cultured for 3 h at 30°C until early exponential phase, followed by induction with L-arabinose (10 mM, Sigma-Aldrich) or 42°C for 20 min, respectively. The cells were washed in ice-cold glycerol (10% v/v) solution and concentrated 100-fold. Ultimately, the PCR product was transformed by electroshock (1.8 V at 5 ms), followed by regeneration in SOC (SOB pre-made mixture, Roth GmbH, and 50 mM D-glucose) medium for 2 h at 37°C, and plated on selective LB-agar plates. The success of the gene knockout was verified by gel electrophoresis and Sanger sequencing (Microsynth AG). Specifically, the multimutant strains Δ4 D-glc and Δ3 D-gal were subjected to whole-genome sequencing to confirm their genetic integrity. Antibiotic resistance cassettes were eliminated via flippase FLP recombination using pCP20 ^58^.

### Homologous recombination by P22 phage transduction

P22 phage transduction was conducted by generating P22 phages containing the antibiotic resistance cassette inserted into the gene of interest from knockout strains generated by λ Red (Supplementary Data 1). The single-gene knockout mutant was incubated overnight with the P22 phage generated from a wild type SB300 background. The culture was treated with chloroform (1% v/v) for 15 min followed by centrifugation and subsequent sterile filtration of the supernatant (0.44 µM pore size). The P22 phages were subsequently incubated with the recipient strain for 15 minutes at 37°C and then plated on selective LB-agar plates. This was followed by two consecutive overnight streaks on selective LB-agar plates. Finally, the transduced clone was examined for P22 phage contamination using Evans Blue Uranine (EBU) LB-agar plates (0.4% w/v glucose, 0.001% w/v Evans Blue, 0.002% w/v Uranine). All mutations were verified by gel electrophoresis or Sanger sequencing (Microsynth AG), using the corresponding primers (Supplementary Data 3).

### Animal experiments

For single and competitive infection experiments, as well as ecological invasion assays, 7 to 12 week old C57BL/6J and 129S6/SvEvTac mice were pretreated with 25 mg streptomycin ^59^. For the LPS-pretreated model, mice were injected intraperitoneally or intravenously with 10 µg ultrapure LPS from *E. coli* O111:B4 (InvivoGen; tlrl-3pelps) in 100 µl PBS ^40^. LPS-treated mice were inoculated with 5 × 10⁷ CFU *Salmonella* at the time of LPS administration, whereas streptomycin-pretreated mice were infected the following day. Mice were inoculated with *S*. Typhimurium cells prepared from an overnight culture grown in selective LB and subcultured for 4 hours at 37°C in LB medium (5% v/v). Bacteria were washed once with PBS, adjusted to an optical density at 600 nm of 2.0, and mice were infected by oral gavage with 5 × 10⁷ CFU. For competitive infection experiments, wild-type and mutant strains were mixed at a 1:1 ratio. For ecological invasion experiments, the wild-type strain was serially diluted 1:1000 in three steps and then combined with the mutant strain. The inoculum of each strain was quantified by plating on selective MacConkey agar and used to normalize bacterial loads recovered from biological samples, including feces, cecal contents, and systemic organs, in the 1:1 competitive infection experiments. The normalized bacterial loads were then used to calculate the competitive index (mutant divided by wild type). For ecological invasion experiments, the 50 µl inoculum was quantified and normalized to CFU per gram of litre. Statistical significance was assessed by comparing the absolute bacterial loads of the wild type and the sugar acid mutant. At the end of the experiments, animals were euthanized by CO₂ asphyxiation. Fresh fecal pellets were collected daily, whereas cecal contents were collected at the end of the experiment. In selected cases, systemic organs including the spleen, liver, and mesenteric lymph nodes were also harvested. All samples were collected daily and homogenized in 500 µl PBS with a metal bead using a TissueLyser (Qiagen). Homogenates were serially diluted and plated on selective MacConkey agar to quantify total bacterial loads.

### Sample preparation for free monosaccharide quantification by LC-MS

Cecum contents were collected in the morning, i.e., the 2nd hour of the light phase in the mouse room, suspended in PBS (1:1 ratio), and homogenized (without metal beads) using a TissueLyser (Qiagen) at 25 Hz for 2 min. This was followed by centrifugation for 5 min at 20,000 g at 4°C to pellet cells and particulate matter. Cell-free supernatants were collected and further centrifuged for 40 min at 20,000 g to remove any remaining suspended particulates. Clear supernatants were then transferred to clean microfuge tubes and stored at -80°C until thawed for LC-MS analysis. The sample (n) size per mouse model was n = 5 biological replicates.

### Quantification of free monosaccharides by LC-MS

Following a previously published protocol ^60^, samples containing free monosaccharides (25 µl) were derivatized with 75 µl of 0.1 M 1-phenyl-3-methyl-5-pyrazolone (PMP) in 2:1 methanol:ddH_2_O with 0.4 % ammonium hydroxide for 100 minutes at 70°C. Additionally, each sample was spiked with an internal standard of 10 µM ^13^C_6_-glucose, ^13^C_6_-galactose and ^13^C_6_-mannose (mass 186 Da). For quantification, we derivatized a serial dilution of a standard mix containing D-mannuronic acid, D-guluronic acid, D-xylose, L-arabinose, D-glucosamine, L-fucose, D-glucose, D-galactose, D-mannose, N-acetyl-D-glucosamine, N-acetyl-D-galactosamine, N-acetyl-D-mannosamine, D-ribose, L-rhamnose and D-galactosamine. Samples and standards were derivatized by incubation at 70°C for 100 minutes. After derivatization, samples were neutralized with HCl, followed by chloroform extraction to remove underivatized PMP, as previously described.

Following ^61^, PMP-derivatives were measured on a SCIEX qTRAP5500 and an Agilent 1290 Infinity II LC system equipped with a Waters CORTECS UPLC C18 Column, 90 Å, 1.6 µm, 2.1 mm X 50 mm reversed phase column with guard column controlled with Sciex Analyst v. 1.7.3. The mobile phase consisted of buffer A (10 mM NH_4_Formate in ddH_2_O, 0.1% formic acid) and buffer B (100% acetonitrile, 0.1% formic acid). PMP-derivatives were separated with an initial isocratic flow of 15% Buffer B for 2 minutes, followed by a gradient from 15% to 20% Buffer B over 5 minutes at a constant flow rate of 0.5 ml/min and a column temperature of 50°C. The ESI source settings were 625°C, with curtain gas set to 30 (arbitrary units), collision gas to medium, ion spray voltage 5500 (arbitrary units), temperature to 625°C, Ion source Gas 1 &2 to 90 (arbitrary units). PMP-derivatives were measured by multiple reaction monitoring (MRM) in positive mode with previously optimized transitions and collision energies. For example, a D-glucose derivative has a Q1 mass of 511 and was fragmented with a collision energy of 35V to yield the quantifier ion of 175 Da and the diagnostic fragment of 217 Da. Different PMP-derivatives were identified by their mass and retention in comparison to known standards. Peak areas were integrated using Skyline 23.1.0.455 and concentrations were quantified using a custom Python v. 3.9. script. In short, technical variation in sample processing were normalized by the amount of internal standard in each sample. Peak areas of the 175 Da fragment were used for quantification using a linear fit to an external standard curve in triplicates with known concentrations ranging from 100 pM to 10 µM.

### Quantification of sugar acids by LC-MS

D-glucose and D-galactose were quantified using the same previously established method as for the other analyzed monosaccharides ^11,18,23^. Carboxyl groups of sugar acids were derivatized with aniline following established protocols ^62,63^. Proteins were precipitated by mixing samples 1:1 (v/v) with methanol containing internal standards (100 µM each of sodium acetate-¹³C₂ (Sigma 282014), succinic acid-d₄ (Sigma 293075), and butyric acid-d₈ (Sigma 588555)). Samples were incubated at −20 °C for 1 h and centrifuged at 10,000 × g for 10 min at 4 °C. For derivatization, 25 µL of supernatant was mixed with 75 µL of derivatization solution (100 mM aniline, 50 mM 1-ethyl-3-(3-dimethylaminopropyl)carbodiimide (EDC)) in water adjusted to pH 4.5 with 50 mM HCl. Samples were incubated at 4 °C for 30 min and the reaction was quenched by addition of 10 µL of 500 mM β-mercaptoethanol. External calibration curves were prepared using a mixture of D-glucuronic acid, D-galacturonic acid, D-glucarate, D-galactarate, D-gluconate, and D-galactonate over a concentration range of 2 µM to 5 mM and processed identically to samples.

Sugar derivatives were measured on a SCIEX qTRAP5500 and an Agilent 1290 Infinity II LC system equipped with a Waters CORTECS UPLC C18 Column, 90 Å, 1.6 µm, 2.1 mm X 50 mm reversed phase column with guard column controlled with Sciex Analyst v. 1.7.3. Chromatographic separation was performed at a flow rate of 500 µL min⁻¹ using a binary solvent system consisting of solvent A (10 mM ammonium formate in water with 0.1% formic acid) and solvent B (acetonitrile with 0.1% formic acid). The gradient was as follows: 0–2.0 min, 100% A; 2.0–7.5 min, linear increase to 30% B; 7.6–8.5 min, 100% B; 8.6–9.5 min, re-equilibration at 100% A. Analytes were identified based on exact mass, characteristic MS/MS fragments, and retention time (Supplementary Data 5). Peak areas were integrated using Skyline 23.1.0.455. Quantification was performed using linear regression of external standards.

### *In vitro* growth experiments

Overnight cultures of *S*. Typhimurium were grown in minimal medium (42 mM Na₂HPO₄, 22 mM KH₂PO₄, 8.6 mM NaCl, 18.7 mM NH₄Cl, supplemented with 2 mM MgSO₄, 0.1 mM CaCl₂, and the indicated carbon source) at 37°C with aeration. Cultures were washed once in sterile PBS and adjusted to the indicated starting optical density (OD_600_). Cells were inoculated into minimal medium supplemented with a single carbon source (20 mM). The following substrates were tested: D-glucose, D-gluconate, D-glucuronate, and D-glucarate, as well as D-galactose, D-galactonate, D-galacturonate, and D-galactarate. Cultures were incubated at 37°C with shaking under aerobic conditions in a 96-well plate layout (150 µl per well). Bacterial growth was monitored by measuring the OD_600_ every 10 minutes for 8 hours using a spectrophotometer. The starting OD_600_ is indicated in the figure. Each condition was tested in independent biological replicates.

### Lipocalin-2 analysis of feces samples

Lipocalin-2 levels were measured in fecal samples homogenized in 500 μl PBS using an ELISA assay (DuoSet Lipocalin-2 ELISA kit, DY1857; R&D Systems) according to the manufacturer’s instructions. Fecal pellets were diluted 1:20, 1:400, or left undiluted, and concentrations were determined using Four-Parametric Logistic Regression curve fitting.

### Haematoxylin and eosin staining of tissue

Haematoxylin and eosin (HE) staining of cryo-embedded tissues (OCT, Sysmex Digitana) and subsequent pathoscoring for granulocyte infiltration were performed as previously described ^59^. The Source Data file includes the individual pathological scores for four subgroups: submucosal edema, epithelial disruption, goblet cell depletion, and neutrophil infiltration. Each score is accompanied by a brief description detailing the different levels of intensity.

### Bioinformatic analysis of sugar acid utilization pathways in *E. coli* from (non-)IBD patient samples

*E. coli* isolates from healthy and IBD patients were obtained from Dubinsky et al., 2022 ^64^. The study collected *E.* coli isolate genome and metagenomic assembled genome data of patients with a pouch, ulcerative colitis (UC), or Crohn’s disease (CD), and from healthy individuals across multiple studies, and additionally generated newly sequenced genomes. Only isolates from healthy individuals and from patients with UC or CD were retained (**Supplementary Data 4**). To assess the presence of D-galactose and D-glucose derived sugar utilization pathways, reference sequences of the respective pathways (see **Fig. 1b** and **Fig. 2a**) were obtained from *E. coli* str. K-12 substr. MG1655 (accession: NC_000913.3). Open reading frames (ORFs) for all selected *E. coli* genomes were predicted using Pyrodigal v3.5.2 ^65^, a Python library binding to Prodigal ^66^. ORFs were deduplicated using CD-HIT v4.8.1^67^ (sequence identity threshold ≥ 95% and alignment coverage ≥ 95%), and a reciprocal best hit search was performed using MMseqs2 ^68^ “easy-rbh” v 18-8cc5c against the reference sequences. Hits with *e*-value > 1e-10, percentage identity < 50% and alignment coverage < 50% were discarded. A sugar acid utilization pathway was considered to be present in a genome, if all respective genes shown in **Fig.1b** and **Fig.2a** were detected (**Supplementary Data 4**). Python 3.7.6 was used to calculate the relative percentage of sugar acid utilization pathways present in *E. coli* genomes from (non-)IBD samples and bar plots were generated using Matplotlib v3.5.3 ^69^.

### Statistical analysis

No statistical methods were used to predetermine sample sizes. For all mouse experiments, sample sizes of 5 or greater were used, consistent with those reported in previous publications ^70^. Data collection and analysis were not performed blind to the conditions of the experiments. Where applicable, two-tailed Mann–Whitney U tests and Wilcoxon rank-sum tests were employed to assess statistical significance, as specified in the figure legends. Statistical analyses were performed using GraphPad Prism 10 for Windows. P values were grouped as follows: **** P < 0.0001; *** P < 0.001; ** P < 0.01; * P < 0.05; ns P ≥ 0.05.

## Notes

### Competing Interest Statement

The authors have declared no competing interest.

## References

1 Deng, L. & Wang, S. Colonization resistance: the role of gut microbiota in preventing Salmonella invasion and infection. Gut Microbes 16, 2424914 (2024). 10.1080/19490976.2024.2424914

2 Caballero-Flores, G., Pickard, J. M. & Nunez, G. Microbiota-mediated colonization resistance: mechanisms and regulation. Nat Rev Microbiol 21, 347–360 (2023). 10.1038/s41579-022-00833-7

3 Herzog, M. K. et al. Mouse models for bacterial enteropathogen infections: insights into the role of colonization resistance. Gut Microbes 15, 2172667 (2023). 10.1080/19490976.2023.2172667

4 Spragge, F. et al. Microbiome diversity protects against pathogens by nutrient blocking. Science 382, eadj3502 (2023). 10.1126/science.adj3502

5 Furuichi, M. et al. Commensal consortia decolonize Enterobacteriaceae via ecological control. Nature 633, 878–886 (2024). 10.1038/s41586-024-07960-6

6 Bakkeren, E., Piskovsky, V., Lee, M. N. Y., Jahn, M. T. & Foster, K. R. Strain displacement in microbiomes via ecological competition. Nature Microbiology (2025). 10.1038/s41564-025-02162-w

7 Eberl, C. et al. E. coli enhance colonization resistance against Salmonella Typhimurium by competing for galactitol, a context-dependent limiting carbon source. Cell Host Microbe 29, 1680–1692 e1687 (2021). 10.1016/j.chom.2021.09.004

8 Wotzka, S. Y., Nguyen, B. D. & Hardt, W. D. Salmonella Typhimurium Diarrhea Reveals Basic Principles of Enteropathogen Infection and Disease-Promoted DNA Exchange. Cell Host Microbe 21, 443–454 (2017). 10.1016/j.chom.2017.03.009

9 Nguyen, B. D. et al. Import of Aspartate and Malate by DcuABC Drives H(2)/Fumarate Respiration to Promote Initial Salmonella Gut-Lumen Colonization in Mice. Cell Host Microbe 27, 922–936 e926 (2020). 10.1016/j.chom.2020.04.013

10 Maier, L. et al. Microbiota-derived hydrogen fuels Salmonella typhimurium invasion of the gut ecosystem. Cell Host Microbe 14, 641–651 (2013). 10.1016/j.chom.2013.11.002

11 Nguyen, B. D. et al. Salmonella Typhimurium screen identifies shifts in mixed-acid fermentation during gut colonization. Cell Host Microbe (2024). 10.1016/j.chom.2024.08.015

12 Winter, S. E. et al. Gut inflammation provides a respiratory electron acceptor for Salmonella. Nature 467, 426–429 (2010). 10.1038/nature09415

13 Winter, S. E. et al. Host-derived nitrate boosts growth of E. coli in the inflamed gut. Science 339, 708–711 (2013). 10.1126/science.1232467

14 Rivera-Chavez, F. et al. Depletion of Butyrate-Producing Clostridia from the Gut Microbiota Drives an Aerobic Luminal Expansion of Salmonella. Cell Host Microbe 19, 443–454 (2016). 10.1016/j.chom.2016.03.004

15 Rogers, A. W. L., Tsolis, R. M. & Baumler, A. J. Salmonella versus the Microbiome. Microbiol Mol Biol Rev 85, 10.1128/mmbr.00027-00019 (2021). 10.1128/MMBR.00027-19

16 Rogers, A. W. L. et al. Salmonella re-engineers the intestinal environment to break colonization resistance in the presence of a compositionally intact microbiota. Cell Host Microbe 32, 1774–1786 e1779 (2024). 10.1016/j.chom.2024.07.025

17 Yoo, W. et al. Salmonella Typhimurium expansion in the inflamed murine gut is dependent on aspartate derived from ROS-mediated microbiota lysis. Cell Host Microbe 32, 887–899 e886 (2024). 10.1016/j.chom.2024.05.001

18 Schubert, C. et al. Monosaccharides drive Salmonella gut colonization in a context-dependent or -independent manner. Nat Commun 16, 1735 (2025). 10.1038/s41467-025-56890-y

19 Schubert, C. & Unden, G. C4-Dicarboxylates as Growth Substrates and Signaling Molecules for Commensal and Pathogenic Enteric Bacteria in Mammalian Intestine. J Bacteriol 204, e0054521 (2022). 10.1128/JB.00545-21

20 Schubert, C. et al. C4-dicarboxylates and l-aspartate utilization by Escherichia coli K-12 in the mouse intestine: l-aspartate as a major substrate for fumarate respiration and as a nitrogen source. Environ Microbiol 23, 2564–2577 (2021). 10.1111/1462-2920.15478

21 Schubert, C., Zedler, S., Strecker, A. & Unden, G. L-Aspartate as a high-quality nitrogen source in Escherichia coli: Regulation of L-aspartase by the nitrogen regulatory system and interaction of L-aspartase with GlnB. Mol Microbiol 115, 526–538 (2021). 10.1111/mmi.14620

22 Schubert, C. et al. Strain-specific galactose utilization by commensal E. coli mitigates Salmonella establishment in the gut. PLoS Pathog 21, e1013232 (2025). 10.1371/journal.ppat.1013232

23 Laganenka, L. et al. Interplay between chemotaxis, quorum sensing, and metabolism regulates Escherichia coli-Salmonella Typhimurium interactions in vivo. PLoS Pathog 21, e1013156 (2025). 10.1371/journal.ppat.1013156

24 Fuchs, L. et al. The Gfr Uptake System Provides a Context-Dependent Fitness Advantage to Salmonella Typhimurium SL 1344 During the Initial Gut Colonization Phase. Molecular Microbiology (2025).

25 Gul, E. et al. Differences in carbon metabolic capacity fuel co-existence and plasmid transfer between Salmonella strains in the mouse gut. Cell Host Microbe 31, 1140–1153 e1143 (2023). 10.1016/j.chom.2023.05.029

26 Ruddle, S. J., Massis, L. M., Cutter, A. C. & Monack, D. M. Salmonella-liberated dietary L-arabinose promotes expansion in superspreaders. Cell Host & Microbe 31, 405–417.e405 (2023). 10.1016/j.chom.2023.01.017

27 Larke, J. A. et al. Dietary Intake of Monosaccharides from Foods is Associated with Characteristics of the Gut Microbiota and Gastrointestinal Inflammation in Healthy US Adults. J Nutr 153, 106–119 (2023). 10.1016/j.tjnut.2022.12.008

28 Castillo, J. J. et al. The Development of the Davis Food Glycopedia-A Glycan Encyclopedia of Food. Nutrients 14 (2022). 10.3390/nu14081639

29 Faber, F. et al. Host-mediated sugar oxidation promotes post-antibiotic pathogen expansion. Nature 534, 697–699 (2016). 10.1038/nature18597

30 Mandrand-Berthelot, M.-A., Condemine, G. & Hugouvieux-Cotte-Pattat, N. Catabolism of hexuronides, hexuronates, aldonates, and aldarates. EcoSal Plus 1, 10.1128/ecosalplus. 1123.1124. 1122 (2004).

31 Proctor, L. M. et al. The Integrative Human Microbiome Project. Nature 569, 641–648 (2019). 10.1038/s41586-019-1238-8

32 Schirmer, M. et al. Dynamics of metatranscription in the inflammatory bowel disease gut microbiome. Nature microbiology 3, 337–346 (2018).

33 Avdagić, N. et al. Nitric oxide as a potential biomarker in inflammatory bowel disease. Bosnian journal of basic medical sciences 13, 5 (2013).

34 Levy, S. et al. Convergent evolution of oxidized sugar metabolism in commensal and pathogenic microbes in the inflamed gut. Nat Commun 16, 1121 (2025). 10.1038/s41467-025-56332-9

35 Näpflin, N., Schubert, C., Malfertheiner, L., Hardt, W.-D. & von Mering, C. Gene-level analysis of core carbohydrate metabolism across the Enterobacteriaceae pan-genome. Communications Biology 8, 1241 (2025). 10.1038/s42003-025-08640-5

36 Boulanger, E. F. et al. Sugar-Phosphate Toxicities Attenuate Salmonella Fitness in the Gut. J Bacteriol 204, e0034422 (2022). 10.1128/jb.00344-22

37 Boulanger, E. F., Sabag-Daigle, A., Thirugnanasambantham, P., Gopalan, V. & Ahmer, B. M. M. Sugar-Phosphate Toxicities. Microbiol Mol Biol Rev 85, e0012321 (2021). 10.1128/MMBR.00123-21

38 Caron, J. et al. Influence of Slc11a1 on the outcome of serovar enteritidis infection in mice is associated with Th polarization. Infection and Immunity 74, 2787–2802 (2006). 10.1128/Iai.74.5.2787-2802.2006

39 Gül, E. et al. The microbiota conditions a gut milieu that selects for wild-type Salmonella Typhimurium virulence. PLoS biology 21, e3002253 (2023).

40 Kroon, S. et al. Sublethal systemic LPS in mice enables gut-luminal pathogens to bloom through oxygen species-mediated microbiota inhibition. Nature Communications 16 (2025). 10.1038/s41467-025-57979-0

41 Shan, Y., Lee, M. & Chang, E. B. The gut microbiome and inflammatory bowel diseases. Annual review of medicine 73, 455–468 (2022).

42 Glassner, K. L., Abraham, B. P. & Quigley, E. M. The microbiome and inflammatory bowel disease. Journal of Allergy and Clinical Immunology 145, 16–27 (2020).

43 Kelleher, Z. T., Matsumoto, A., Stamler, J. S. & Marshall, H. E. NOS2 regulation of NF-κB by S-nitrosylation of p65. Journal of biological chemistry 282, 30667–30672 (2007).

44 Moreira de Gouveia, M. I., Bernalier-Donadille, A. & Jubelin, G. Enterobacteriaceae in the human gut: dynamics and ecological roles in health and disease. Biology 13, 142 (2024).

45 Mirsepasi-Lauridsen, H. C., Vallance, B. A., Krogfelt, K. A. & Petersen, A. M. Escherichia coli pathobionts associated with inflammatory bowel disease. Clinical microbiology reviews 32, 10.1128/cmr.00060-00018 (2019).

46 Doranga, S., Krogfelt, K. A., Cohen, P. S. & Conway, T. Nutrition of Escherichia coli within the intestinal microbiome. EcoSal Plus 12, eesp00062023 (2024). 10.1128/ecosalplus.esp-0006-2023

47 Doranga, S. & Conway, T. Nitrogen assimilation by E. coli in the mammalian intestine. mBio 15, e0002524 (2024). 10.1128/mbio.00025-24

48 Jones, S. A. et al. Anaerobic Respiration of Escherichia coli in the Mouse Intestine. Infection and Immunity 79, 4218–4226 (2011). 10.1128/iai.05395-11

49 Jones, S. A. et al. Respiration of Escherichia coli in the mouse intestine. Infect Immun 75, 4891–4899 (2007). 10.1128/IAI.00484-07

50 Yip, A. Y. G. et al. Antibiotics promote intestinal growth of carbapenem-resistant Enterobacteriaceae by enriching nutrients and depleting microbial metabolites. Nat Commun 14, 5094 (2023). 10.1038/s41467-023-40872-z

51 Porter, N. T. & Martens, E. C. The Critical Roles of Polysaccharides in Gut Microbial Ecology and Physiology. Annual Review of Microbiology 71, 349–369 (2017). 10.1146/annurev-micro-102215-095316

52 Morgan, X. C. et al. Dysfunction of the intestinal microbiome in inflammatory bowel disease and treatment. Genome biology 13, R79 (2012).

53 Lewis, J. D. et al. Inflammation, antibiotics, and diet as environmental stressors of the gut microbiome in pediatric Crohn’s disease. Cell host & microbe 18, 489–500 (2015).

54 Santana, P. T., Rosas, S. L. B., Ribeiro, B. E., Marinho, Y. & de Souza, H. S. Dysbiosis in inflammatory bowel disease: pathogenic role and potential therapeutic targets. International journal of molecular sciences 23, 3464 (2022).

55 Hall, A. B. et al. A novel Ruminococcus gnavus clade enriched in inflammatory bowel disease patients. Genome medicine 9, 103 (2017).

56 Hug, L. A. et al. A roadmap for equitable reuse of public microbiome data. Nature microbiology, 1–12 (2025).

57 Datsenko, K. A. & Wanner, B. L. One-step inactivation of chromosomal genes in Escherichia coli K-12 using PCR products. Proceedings of the National Academy of Sciences 97, 6640–6645 (2000).

58 Cherepanov, P. P. & Wackernagel, W. Gene disruption in Escherichia coli: TcR and KmR cassettes with the option of Flp-catalyzed excision of the antibiotic-resistance determinant. Gene 158, 9–14 (1995).

59 Barthel, M. et al. Pretreatment of Mice with Streptomycin Provides a *Salmonella enterica* Serovar Typhimurium Colitis Model That Allows Analysis of Both Pathogen and Host. Infection and Immunity 71, 2839–2858 (2003). 10.1128/iai.71.5.2839-2858.2003

60 Ruhmann, B., Schmid, J. & Sieber, V. Fast carbohydrate analysis via liquid chromatography coupled with ultra violet and electrospray ionization ion trap detection in 96-well format. J Chromatogr A 1350, 44–50 (2014). 10.1016/j.chroma.2014.05.014

61 Xu, G., Amicucci, M. J., Cheng, Z., Galermo, A. G. & Lebrilla, C. B. Revisiting monosaccharide analysis - quantitation of a comprehensive set of monosaccharides using dynamic multiple reaction monitoring. Analyst 143, 200–207 (2017). 10.1039/c7an01530e

62 Meng, X. et al. Simultaneous 3-nitrophenylhydrazine derivatization strategy of carbonyl, carboxyl and phosphoryl submetabolome for LC-MS/MS-based targeted metabolomics with improved sensitivity and coverage. Analytical chemistry 93, 10075–10083 (2021).

63 Chan, J. C. Y., Kioh, D. Y. Q., Yap, G. C., Lee, B. W. & Chan, E. C. Y. A novel LCMSMS method for quantitative measurement of short-chain fatty acids in human stool derivatized with 12C-and 13C-labelled aniline. Journal of pharmaceutical and biomedical analysis 138, 43–53 (2017).

64 Dubinsky, V. et al. Escherichia coli strains from patients with inflammatory bowel diseases have disease-specific genomic adaptations. Journal of Crohn’s and Colitis 16, 1584–1597 (2022).

65 Larralde, M. Pyrodigal: Python bindings and interface to Prodigal, an efficient method for gene prediction in prokaryotes. Journal of Open Source Software 7, 4296 (2022).

66 Hyatt, D. et al. Prodigal: prokaryotic gene recognition and translation initiation site identification. BMC bioinformatics 11, 119 (2010).

67 Li, W. & Godzik, A. Cd-hit: a fast program for clustering and comparing large sets of protein or nucleotide sequences. Bioinformatics 22, 1658–1659 (2006).

68 Steinegger, M. & Söding, J. MMseqs2 enables sensitive protein sequence searching for the analysis of massive data sets. Nature biotechnology 35, 1026–1028 (2017).

69 Hunter, J. D. Matplotlib: A 2D graphics environment. Computing in science & engineering 9, 90–95 (2007).

70 Stecher, B. r., et al. Flagella and chemotaxis are required for efficient induction of Salmonella enterica serovar Typhimurium colitis in streptomycin-pretreated mice. Infection and immunity 72, 4138–4150 (2004).

