## Supplementary Figures 1-7 for "D-gluconate drives *Salmonella* growth during acute and chronic infection"

### Supplementary Figure 1

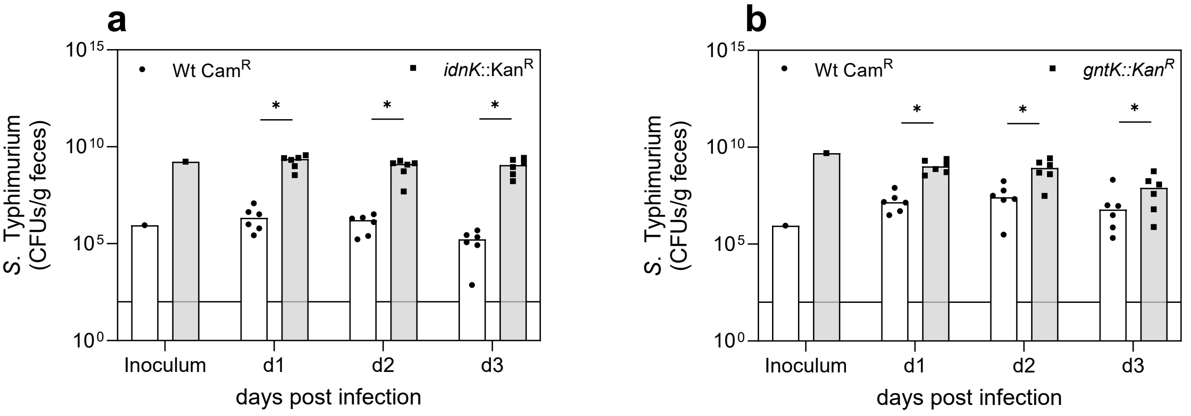


**Supplementary Figure 1. Individual contributions of D-gluconate phosphorylation to wild-type *S.* Typhimurium expansion *in vivo*. a,b,** Fecal colonization levels of *S.* Typhimurium wild type and mutants lacking *idnK* (**a**) or *gntK* (**b**) following oral infection of streptomycin-pretreated C57BL/6J mice. Bacterial loads are shown as CFU per gram feces at the indicated days post infection. The inoculum was normalized to CFU per gram. Each dot represents an individual mouse; bars indicate median values. Statistical significance was assessed using the non-parametric Wilcoxon rank-sum test; p-values are indicated (*: $p < 0.05$).

### Supplementary Figure 2

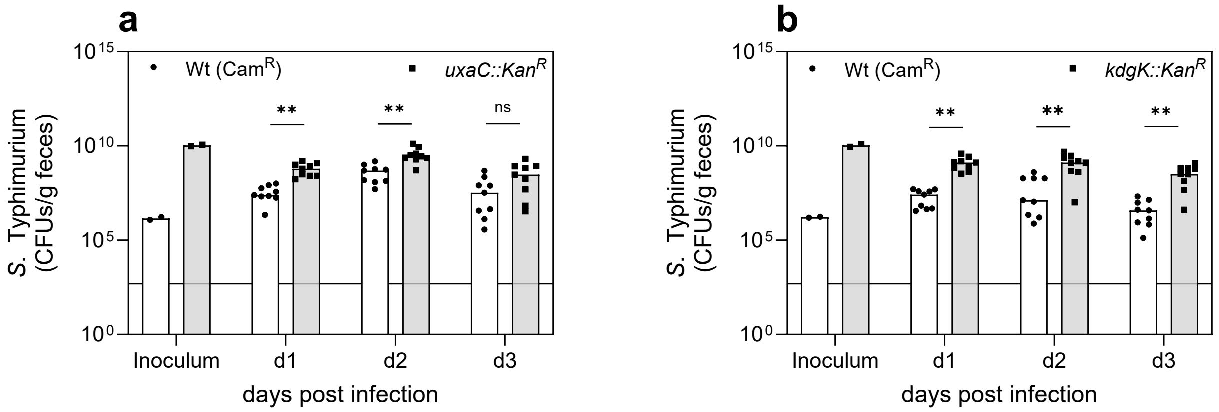


**Supplementary Figure 2. Contribution of uronic acid catabolic steps and KdgK to *S*. Typhimurium wild type expansion. a,b**, Fecal colonization levels of *S*. Typhimurium wild type ($Cam^R$) and mutants lacking *uxaC* (**a**) or *kdgK* (**b**) following oral infection of streptomycin-pretreated C57BL/6J mice. Bacterial loads are shown as CFU per gram feces at the indicated days post infection. The inoculum was normalized to CFU per gram. Each dot represents an individual mouse; bars indicate median values. Statistical significance was assessed using the non-parametric Wilcoxon rank-sum test; p values are indicated (not significant, ns: $p \geq 0.05$; **: $p < 0.01$).

### Supplementary Figure 3

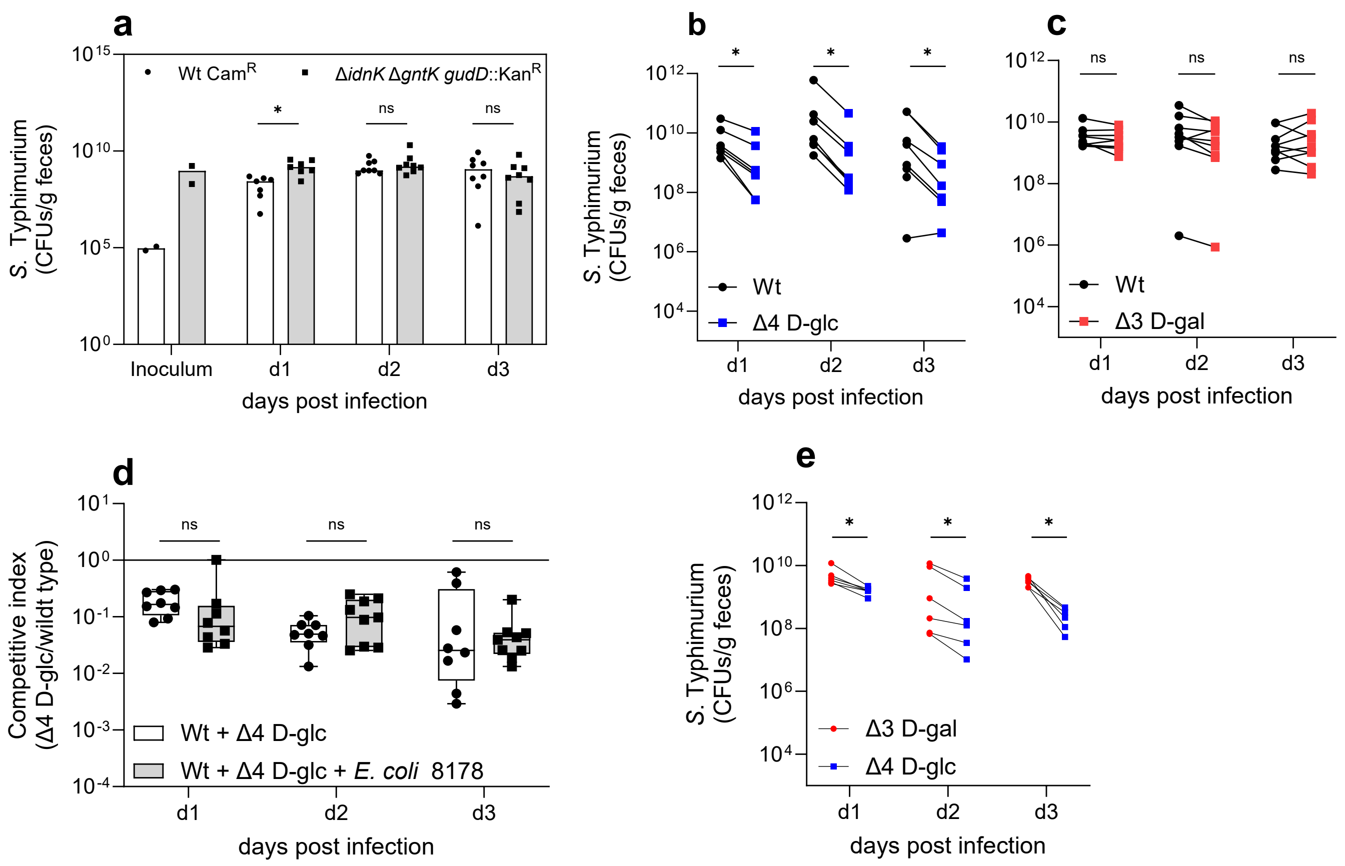


**Supplementary Figure 3. Loss of D-glucose derived sugar acid utilization affects competitive fitness. a,** Fecal colonization levels of *S.* Typhimurium wild type (Cam$^R$) and a mutant lacking D-gluconate and D-glucarate utilization pathways (Δ*idnK* Δ*gntK* Δ*gudD*) following oral infection of streptomycin-pretreated C57BL/6J mice. Bacterial loads are shown as CFU per gram feces at the indicated days post infection. **b,c,** Paired fecal bacterial loads of wild type and mutants deficient in D-glucose-derived (Δ4 D-glc, **b**) or D-galactose-derived (Δ3 D-gal, **c**) sugar acid utilization measured in individual mice over time. Lines connect paired samples from the same animal. **d**, Competitive indices (Δ4 D-glc vs. wild type) were determined in the streptomycin-pretreated mouse model over the course of infection (days 1–3 post-infection). Mice were infected with wild type and Δ4 D-glc either alone or together with *E. coli* 8178, a previously described nutrient competitor of *S.* Typhimurium. Competitive indices are shown on a logarithmic scale. No significant differences (ns) were observed between conditions at any time point, indicating that the presence of *E. coli* 8178 does not alter the fitness of the Δ4 D-glc mutant. **e,** Direct comparison of fecal bacterial loads of Δ3 D-gal and Δ4 D-glc mutants within the same animals, highlighting reduced fitness of the Δ4 D-glc strain. Statistical significance was assessed using the non-parametric Wilcoxon rank-sum test for subpanels **a**, **b**, **c** and **e**, and a two-tailed Mann–Whitney U test for subpanel d; p-values are indicated (not significant, ns: $p \geq 0.05$; *: $p < 0.05$; **: $p < 0.01$; ***: $p < 0.001$)

### Supplementary Figure 4

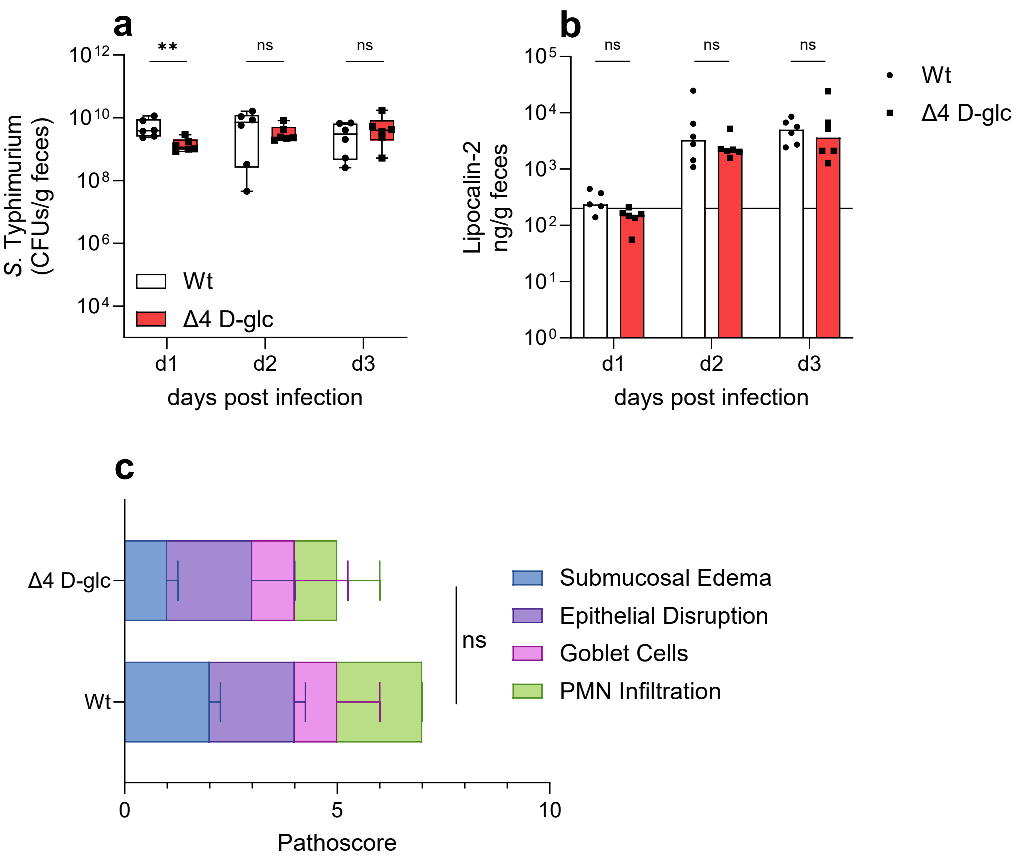


**Supplementary Figure 4. Loss of D-glucose derived sugar acid utilization does not alter inflammatory pathology. a,** Fecal colonization levels of wild type and Δ4 D-glc mutant following single-strain infection, indicating no major defect in absolute colonization capacity. **b,** Fecal lipocalin-2 concentrations at the indicated days post infection, showing comparable levels of intestinal inflammation between wild type and Δ4 D-glc infections. **c,** Histopathological scores of cecal tissue collected at day 3 post infection, including submucosal edema, epithelial disruption, goblet cell depletion, and polymorphonuclear neutrophil (PMN) infiltration, demonstrating no significant differences between wild type and Δ4 D-glc infection. Each dot represents an individual mouse; bars or box plots indicate median values. Statistical significance was assessed using a two-tailed Mann–Whitney U test; *P* values are indicated (not significant, ns: $p \geq 0.05$; **: $p < 0.01$)

### Supplementary Figure 5

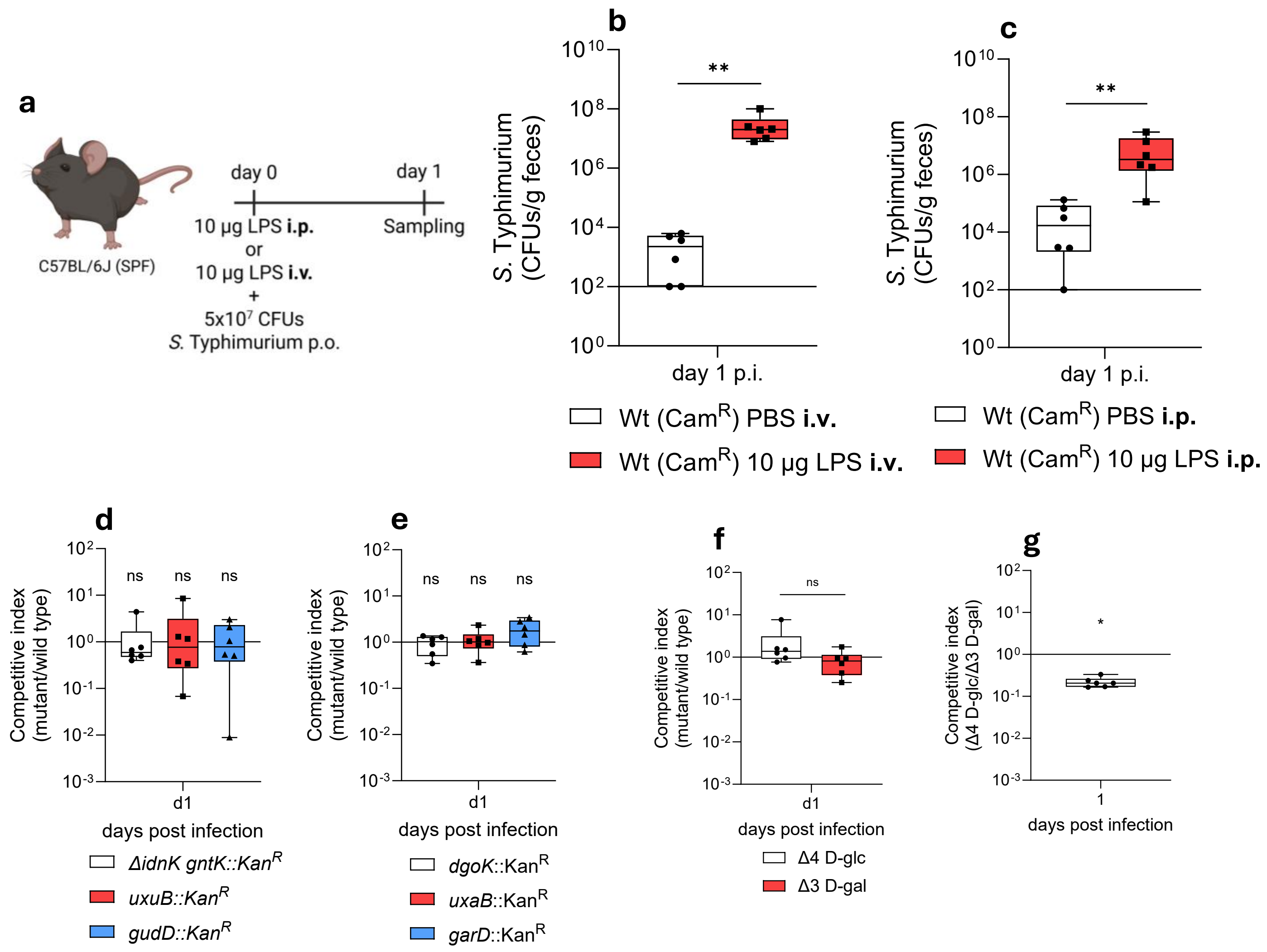


**Supplementary Figure 5. Lipopolysaccharide (LPS)-triggerd ROS promotes *S.* Typhimurium expansion but does not confer a competitive advantage via sugar acid utilization. a,** Experimental design of the LPS mouse model. C57BL/6J specific-pathogen-free mice were treated intraperitoneally or intravenously with lipopolysaccharide (LPS; 10 µg) at the time of oral infection with 5 × $10^7$ CFU of *S.* Typhimurium. Fecal samples were collected one day post infection. Created in BioRender. Schubert, C. (2026) https://BioRender.com/3p40qc2. **b,c,** Fecal colonization levels of *S.* Typhimurium wild type (Cam$^R$) one day post infection following treatment with PBS or LPS, demonstrating increased bacterial loads in LPS-treated mice. **d,e,** Competitive index of wild type versus mutants deficient in D-glucose-derived sugar acid utilization pathways including D-gluconate (Δ*idnK* Δ*gntK*), D-glucuronate (Δ*uxuB*), and D-glucarate (Δ*gudD*) (**d**), or D-galactose-derived sugar acid utilization pathways including D-galactonate (Δ*dgoK*), D-galacturonate (Δ*uxaB*), and D-galactarate (Δ*garD*) (**e**), one day post infection, indicating no significant competitive advantage in the LPS mouse model. **f,** Competitive index of mutants lacking combined Δ4 D-glc and Δ3 D-gal utilization pathways relative to wild type one day post infection. **g,** Competitive index comparing Δ4 D-glc and Δ3 D-gal sugar acid utilization-deficient strains, showing attenuated fitness of Δ4 D-glc in the LPS model. Each dot represents an individual mouse; bars or box plots indicate median values. Statistical significance was assessed using the non-parametric Wilcoxon rank-sum test for subpanels d, e, and g, and a two-tailed Mann–Whitney U test for subpanel b, c, and f; p values are indicated (not significant, ns: $p \geq 0.05$; *: $p < 0.05$; **: $p < 0.01$)

### Supplementary Figure 6

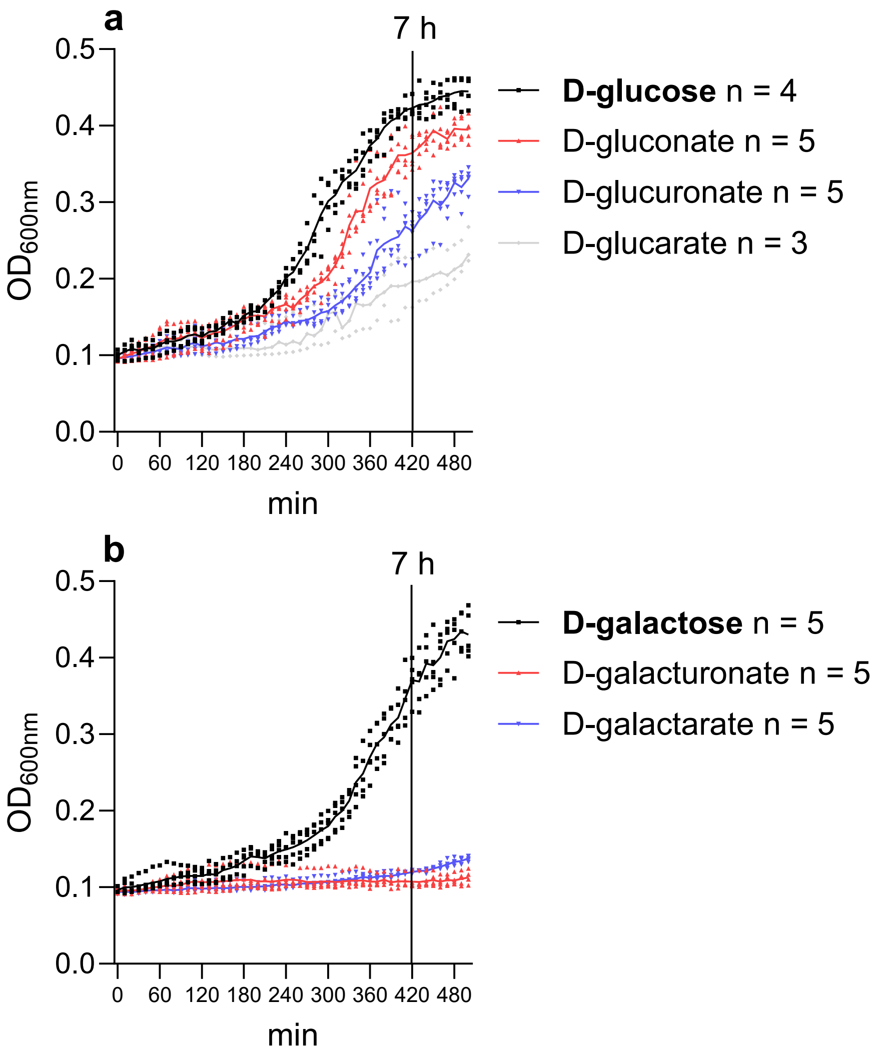


**Supplementary Figure 6. Growth curves of *S.* Typhimurium on sugar acids *in vitro*.**
**a,b,** Growth curves of *S.* Typhimurium cultured in minimal medium supplemented with D-glucose or its derived sugar acids (**a**) and D-galactose or its derived sugar acids (**b**). Optical density ($OD_{600}$) was monitored over time; the 7 h time point used for endpoint analyses is indicated. Data represent the mean of n = 4 (**a**) or n = 5 (**b**) independent biological replicates.

### Supplementary Figure 7

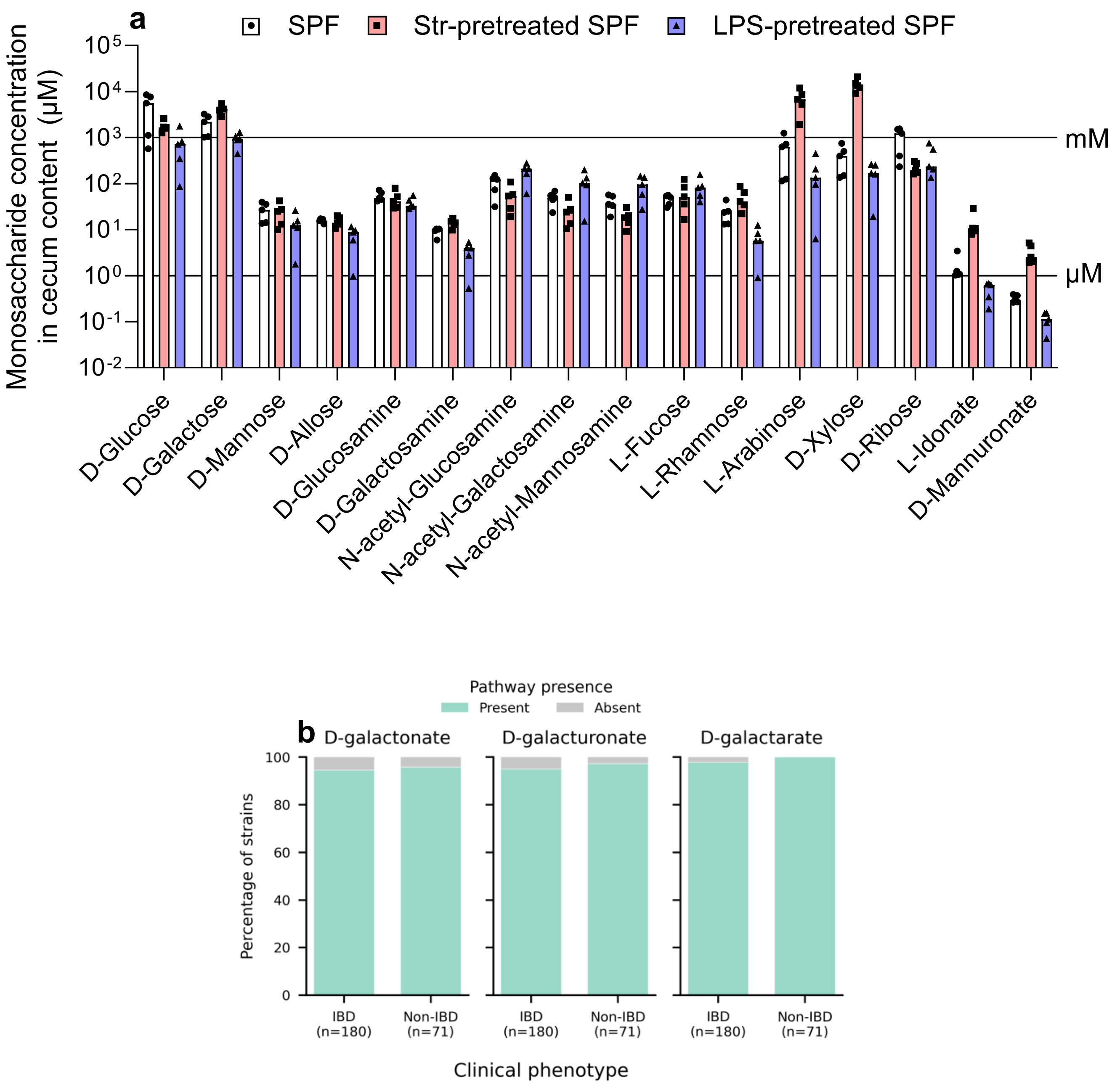


**Supplementary Figure 7. Sugar acids are present in the murine cecum at physiologically relevant concentrations under different inflammatory conditions. a,** Concentrations of monosaccharides in cecal contents of specific-pathogen-free (SPF), streptomycin-pretreated SPF, and LPS-pretreated C57BL/6J SPF mice. Data are displayed on a logarithmic scale; horizontal reference lines indicate micromolar (µM) and millimolar (mM) concentration ranges. Each symbol represents an individual mouse; bars indicate median values. **b**, Bar plots show the percentage of *E. coli* strains harboring the indicated metabolic pathways in isolates derived from patients with inflammatory bowel disease (IBD, n = 180) and non-IBD controls (n = 71). Pathway presence was determined based on genomic analysis. For all three pathways, the vast majority of strains in both cohorts encode the complete degradation pathways, with only a small fraction lacking the respective genes. No substantial differences in pathway prevalence were observed between IBD and non-IBD-derived isolates.
